# Graph-based Contrastive Learning Enables Unified Integration and Niche Transfer Across Single-Cell and Spatial Multi-Omics

**DOI:** 10.1101/2025.02.19.638965

**Authors:** Weige Zhou, Xueying Fan, Lanxiang Li, Jianrong Zheng, Xiaodong Liu, Wenfei Jin, Luyi Tian

## Abstract

The rapid growth of single-cell and spatial omics has outpaced computational methods capable of unifying these data into a cohesive framework for tissue atlas construction and cross-sample analysis. A critical bottleneck lies in the inability of existing tools to co-embed cells from diverse technologies—spanning transcriptomics, epigenomics, and proteomics—into a shared reference space while preserving spatial architecture and molecular specificity. Here, we present Garfield (Graph-based Contrastive Learning Enables Fast Single-Cell Embedding), a geometric deep-learning framework that addresses these challenges through spatially or molecularly aware cell embedding. Leveraging a graph contrastive learning framework, Garfield learns a shared embedding space for data generated by diverse technologies, enabling seamless construction and querying of spatial reference atlases. Our results show that Garfield consistently outperforms state-of-the-art benchmark models in identifying spatial niches across multiple datasets. We further demonstrate Garfield’s versatility by applying it to multi-modal spatial data, including gene expression and chromatin accessibility, where it successfully identifies distinct niches in the mouse brain. Notably, Garfield reveals tumor microenvironment heterogeneity in non-small cell lung cancer and breast cancer, uncovered conserved, barrier-like immune niches at tumor margins orchestrating CD80-mediated T cell–B cell–dendritic cell interactions and IFN-/B cell activation pathways, forming spatially coordinated immune surveillance hubs. These findings underscore Garfield’s potential to advance spatial omics research by offering a robust, scalable solution for integrating and interpreting complex spatial data across diverse tissue types and modalities.

## 1 Introduction

Recent advances in single-cell omics have enabled both individual and integrated cellular profiling. Single-cell multi-omics technologies now enable simultaneous measurements across multiple cellular layers, such as transcriptomics, epigenomics, and proteomics, significantly enhancing our understanding of cell states and the molecular mechanisms underlying development and disease. For example, CITE-seq [1] and REAP-seq [2] enable simultaneous detection of RNA expression and surface protein abundance in single cells, while SNARE-seq [3], SHARE-seq [4], 10x Multiome (https://www.10xgenomics.com/products/) and ISSAAC-seq [5] provide information on both RNA expression and chromatin accessibility from the same cell. Building on these advancements, spatial transcriptomics has emerged as a powerful approach that maps gene expression within the spatial context of tissues, adding a new layer of information. With the expansion to spatial multi-omics, researchers can now profile multiple omic layers (e.g., transcriptomics, epigenomics) within a single tissue section simultaneously, capturing complementary biological insights. Current spatial technologies are broadly divided into sequencing-based [6–10], which infer spatial information by capturing and sequencing RNA, and imaging-based [11– 13] approaches, which directly visualize RNA distribution using microscopy, each unlocking multi-dimensional, spatially-resolved perspectives of each cell.

Despite their transformative potential, computational challenges persist in maximizing the utility of these technologies. Integration methods for single-cell multimodal data often depend on identifying “anchors” to align datasets, either explicitly or implicitly. Based on anchor choice, integration approaches are typically categorized into horizontal, vertical, and diagonal integration [14]. We focus on state-of-the-art algorithms for horizontal and vertical integration. Horizontal integration, as in Seurat v3’s CCA [15], MultiVI [16], Multigrate [17], and MOFA+ [18], uses shared modalities to link datasets with differing cells. Conversely, vertical integration aligns datasets across omics layers, illustrated by tools like Seurat v4’s WNN [19] and MultiVI [16]. Spatial computational methods for identifying cell niches, such as GraphST [20], STAGATE [21], CellCharter [22], SPACE [23] and NicheCompass [24], have also been developed, yet these are designed solely for spatial transcriptomic data. Consequently, integrating or analyzing single-cell or spatial transcriptome data often requires multiple tools, increasing the user’s workload. Furthermore, current methods largely lack functionality to fully leverage relationships across cellular features and multi-modalities within a single cell. Limited tools supports joint analysis of single-cell and spatial multi-omics data while preserving both molecular specificity and spatial topology—a critical requirement for constructing comprehensive tissue atlases.

To address these limitations, we present Garfield (Graph-based Contrastive Learning enables Fast Single-Cell Embedding), a versatile embedding framework that unifies heterogeneous single-cell and spatial (multi-omics) datasets within a shared latent space, supporting a variety of analysis tasks. Unlike existing methods designed specifically for either single-cell or spatial data, Garfield simultaneously processes both data types, regardless of modality. Garfield constructs a unified cell graph where nodes represent individual cells and edges define inter-cellular relationships. By leveraging Variational Graph Autoencoders (VGAE) [25] alongside a contrastive strategy based on SVD and optional MMD-based alignment, Garfield embeds cells into a low-dimensional, batch-invariant biological space, enabling batch correction and multiomics integration at the single-cell or spatial levels. Inspired by architectural surgery [26], Garfield further enables weight-constrained fine-tuning to map new datasets to single-cell or spatial reference atlases.

Garfield’s unified framework supports tasks including: (1) dimensionality reduction; (2) batch correction and multi-modal integration; (3) niche discovery in spatial data; and (4) QueryToReference mapping. Its adaptability across tasks is achieved by adjusting the input graph constructed from single-cell or spatial data. Extensive testing on multiple single-cell multi-omics, spatial uni-modal, and dual-omics datasets demonstrates Garfield’s robust performance, either surpassing or matching current state-of-the-art methods. Additionally, Garfield efficiently scales to large spatial datasets, enabling the construction of comprehensive atlases spanning nearly a million cells through mini-batch processing. Finally, we have developed a comprehensive, scalable Python package with a seamless workflow for data preprocessing, graph construction, graph embedding training with PyTorch, and post-analysis visualization. Compatible with popular single-cell (Scanpy)[27] and spatial tools (Squidpy)[28], Garfield provides detailed documentation and tutorials available at https://garfield-bio.readthedocs.io.

## 2 Results

### 2.1 Overview of Garfield

Garfield is a novel and versatility embedding algorithm designed to support high-resolution analyses of single-cell or spatial data in both uni- and multi-modality formats. By integrating multi-omics with optional spatial information, Garfield deciphers complex molecular landscapes and spatial niches within tissue samples (if provided spatial information). Its flexible input includes feature matrices of segmented cells or capture locations (such as beads, pixels, bins, or spots), along with spatial coordinates. To streamline description, we refer to both cells and capture locations as “cells”, avoiding restriction to any specific platform or resolution.

Garfield employs a Variational Graph Autoencoder (VGAE) [25] framework to embed cells within a spatially and molecularly coherent latent space using a self-supervised, multi-task learning approach. For single-cell data, Garfield constructs a molecular neighborhood graph from single-cell omics datasets, with nodes representing cells and edges indicating cell similarity at the molecular levels. For spatial data, it builds a spatial neighborhood graph using cell or spot coordinates, where edges capture spatial relationships. Each node encodes a feature vector representing omics data, such as gene expression in unimodal analyses or paired gene expression with either chromatin accessibility peaks or antibodies in multimodal setups. Unlike previous methods [23, 29] that rely on single graph type (either molecular graph or spatial graph), Garfield concurrently integrates intra-modality molecular graph, inter-modality molecular graph, and spatial graph if available (derived from spatial coordinates).

To achieve spatially and molecularly integrated latent representations, Garfield offers several graph neural network (GNN) encoders (GATv2 [30], GAT [31], and GCN [32]). To enhance biological signal while minimizing noise, Garfield incorporates a sparse approximation of Singular Value Decomposition (approxSVD) contrastive learning [33] strategy, maximizing mutual information between original and denoised node views. Unlike traditional variational autoencoders that rely on the encoder to estimate both data mean and variance, Garfield’s encoder learns only the mean, with the variance calculated based on this mean through an additional fully connected layer. This adaptive variance scaling adjusts contrast loss per node, inherently reflecting biological variation across cells. Additionally, to mitigate batch effects, Garfield applies Maximum Mean Discrepancy (MMD) [34] loss as an alignment strategy, which is widely utilized in batch correction [26, 35].

In the decoding phase, Garfield reconstructs both spatial and molecular information via multiple modules. A graph decoder aligns sample-specific latent representations to reconstruct spatial graphs through edge reconstruction loss, preserving spatial structure by enforcing similarity between neighboring nodes. For molecular information, Garfield utilizes an omics decoder composed of an inner product module and a feature reconstruction module to optimize omics feature recovery.

In essence, Garfield’s comprehensive framework is an expanded multimodal graph variational autoencoder. This design enables precise characterization of single-cell landscapes and tissue-specific niches, offering a versatile, end-to-end framework for single-cell and spatial omics data analysis.

### 2.2 Benchmarking Garfield for Single-Cell Multi-Omics Integration

Single-cell multi-omics technologies enable researchers to integrate genomic information, providing insights into cellular functions[19], gene regulation[36], and precise fate prediction[37, 38]. To evaluate Garfield’s performance in single-cell multi-omics integration, we compared it with four widely used algorithms—Seurat V4, MultiVI, MOFA+, and Multigrate—on nine scRNA + scATAC datasets (Fig. 2a-c, Supplementary Table 1). Clustering was performed using the Leiden algorithm, matching the number of clusters to ground truth cell types. Performance was assessed with four metrics: adjusted Rand index (ARI), normalized mutual information (NMI), cell-type average silhouette width (cASW), and cell-type separation local inverse Simpson’s index (cLISI). The biological variation conservation score, calculated as the average of these metrics, was used as an overall performance indicator.

**Fig. 1.**
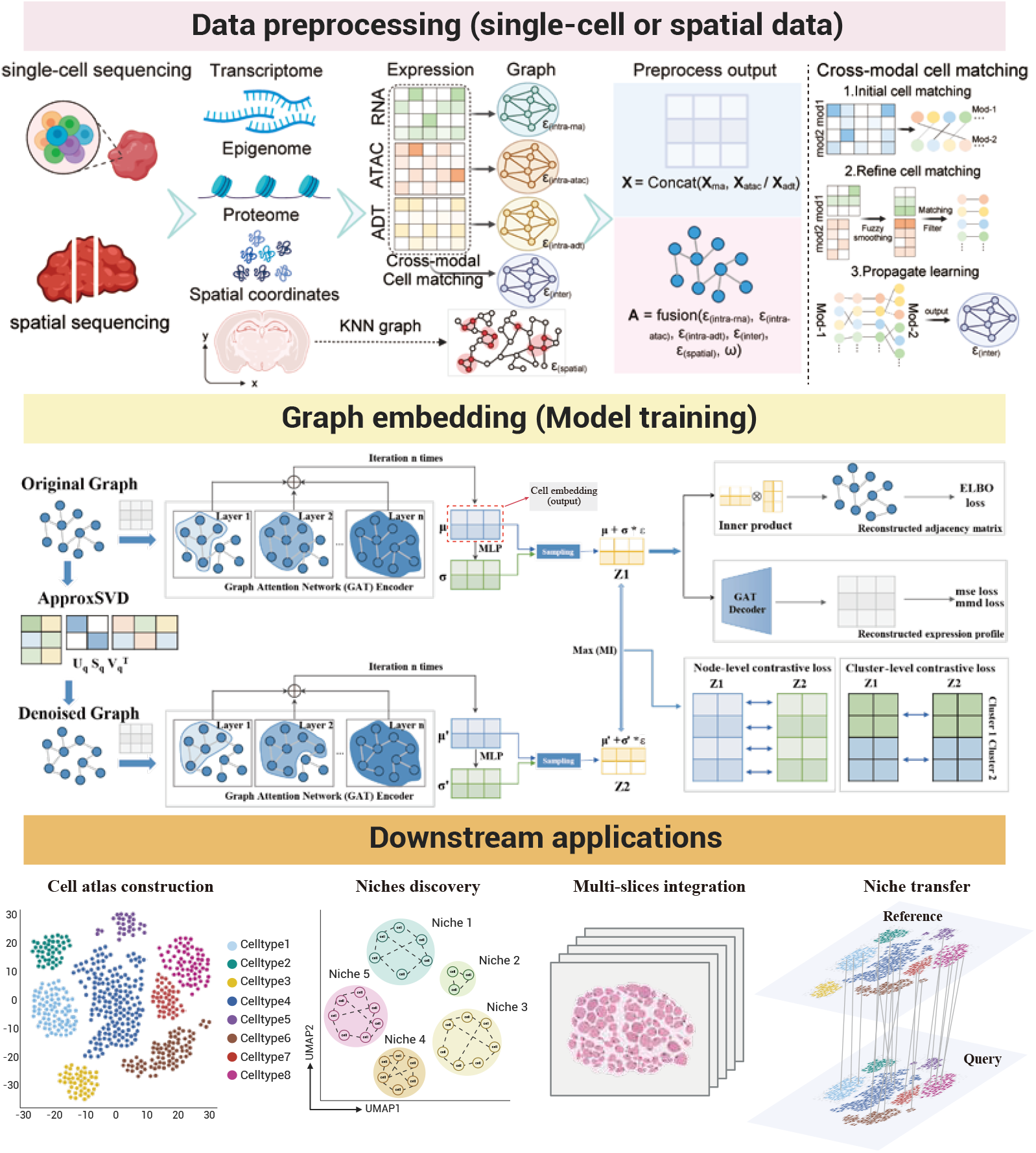
Garfield framework overview. Garfield takes single- or multi-modal single-cell and spatial omics data as input, embedding cells within a shared latent space to address core tasks in single-cell and spatial data analysis. Top: Examples of biological entities that Garfield can encode include the transcriptome, epigenome, proteome, and spatial omics. Middle: Using a graph neural network encoder, Garfield encodes input omics features into a latent feature space, iteratively aggregating neighbor representations. Aided by contrastive learning and the Variational Graph Autoencoder (VGAE) architecture, this representation becomes both batch-invariant and spatially-aware, capturing the complexity of biological space. Bottom: Garfield empowers essential downstream applications in single-cell and spatial omics analysis. Figure created with BioRender.com.

**Fig. 2.**
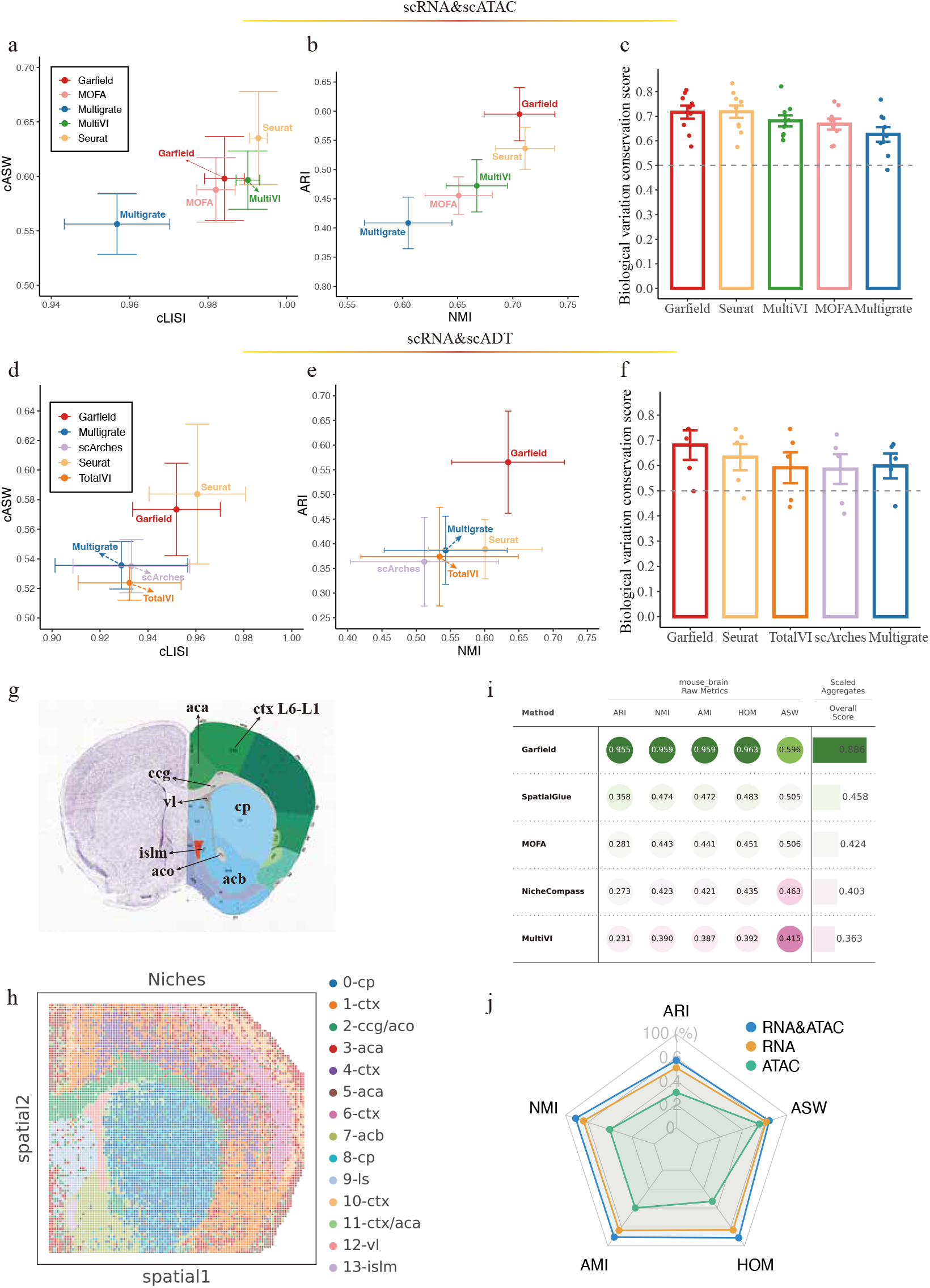
Garfield outperforms existing state-of-the-art methods in single-cell or spatial multi-modal integration. **a**. The average ARI and NMI values of five vertical integration algorithms combining RNA expression and chromatin data across nine single-cell RNA + ATAC datasets. The x and y axes represent the average cLISI and cASW values, respectively. Error bars denote the standard error (s.e.) across the 9 datasets, with data presented as mean ± s.e. **b**. The same analysis as in panel a, but results are evaluated using the average NMI and ARI values. **c**. Bar plots displaying the overall performance of these algorithms, evaluated by biological variation conservation scores across 9 paired RNA + ATAC datasets. Data are shown as mean values with 95% confidence intervals (N = 9 datasets). Each dot represents the biological variation conservation score for an algorithm applied to a specific dataset. **d–f**. Similar to panels a–c, but the results were obtained using five algorithms that integrate RNA expression and protein abundance across five paired RNA + protein datasets. Data are presented as mean values with 95% confidence intervals (N = 5 datasets) **g**. Annotated coronal section of the mouse brain, derived from the Allen Mouse Brain Atlas, serving as a reference for anatomical regions. **h**. Spatial map showing the distribution of Garfield-derived niches, with each niche represented by a distinct color. This result will serve as the pseudo-ground truth in subsequent benchmarking analysis. cp, caudoputamen; ctx, cerebral cortex; ccg, genu of corpus callosum; aco, anterior commissure (olfactory limb); aca, anterior cingulate area; acb, nucleus accumbens; ls, lateral septal nucleus; vl, lateral ventricle; lpo, lateral preoptic area; islm, major island of calleja. **i**. The complex table plot summarizing performance metrics (ARI, NMI, AMI, HOM, ASW) across multiple methods, including Garfield, SpatialGlue, MOFA+, NicheCompass, and MultiVI. **j**. Radar chart comparing analytical performance for RNA, ATAC, and combined RNA+ATAC modalities using five metrics.

Garfield achieved the highest scores, including an average ARI of 0.59, NMI of 0.71, cASW of 0.59, and cLISI of 0.98 (Fig. 2a,b). Seurat V4 also performed well with ARI of 0.53, NMI of 0.71, cASW of 0.63, and cLISI of 0.99, resulting in biological variation conservation scores of 0.716 and 0.718 for Garfield and Seurat V4, respectively, outperforming MultiVI (0.681), Multigrate (0.626), and MOFA+ (0.668; Fig. 2c). Garfield was further evaluated alongside Seurat V4, TotalVI, scArches, and Multi-grate on five scRNA + scADT datasets (Supplementary Table 2). Garfield displayed the highest average ARI (0.57), NMI (0.63; Fig. 2e), and superior biological variation conservation (Garfield: 0.68; Seurat V4: 0.63; TotalVI: 0.59; scArches: 0.59; Multi-grate: 0.60; Fig. 2f). The detailed performance metrics of various algorithms across different datasets are presented in Fig. S1. For clarity, we highlight one representative dataset that showcases Garfield’s ability to capture cross-modality representations from parallel single-cell RNA–ATAC or RNA–ADT datasets (Fig. S2–S3).

Overall, comprehensive benchmarking shows Garfield outperforms or matches current mainstream tools in integrating single-cell multi-omics (scRNA + scATAC and scRNA + scADT).

### 2.3 Benchmarking Garfield for Spatial Multi-Omics Integration

Spatial transcriptomics represents the next frontier in biological sample analysis following the success of single-cell transcriptomics[39]. With advancements in spatial technologies, it is now possible to profile multiple omics layers simultaneously on a single tissue section[40]. Notably, Garfield is also suitable for integrating spatial multi-omics data.

To evaluate Garfield’s capabilities, we benchmarked it against competing methods—NicheCompass, SpatialGlue, MultiVI, and MOFA+—using a mouse brain epigenome–transcriptome dataset. The Allen Brain Atlas reference was used to annotate anatomical regions such as the anterior cingulate area (aca), cortex layers (ctx), genu of corpus callosum (ccg), lateral septal nucleus (ls), major island of Calleja (islm), and nucleus accumbens (acb; Fig. 2g-h). Distinct spatial distributions were observed for each anatomical region, with clear conformity between subcomponents (Fig. S4a-b).

Benchmarking involved applying Leiden clustering across various resolutions on the integrated results from each method to minimize bias from resolution parameters (Methods). Performance in spatial niche detection was assessed using five metrics: adjusted Rand index (ARI), normalized mutual information (NMI), adjusted mutual information (AMI), homogeneity (Homo), and niche average silhouette width (ASW). Garfield outperformed competitors, achieving improvements of 59.7%, 48.5%, 48.7%, 48.0%, and 9.0% over the second-best method across these metrics, with an average improvement of 42.78% (Fig. 2i).

Qualitative evaluation through UMAP and spatial visualizations demonstrated Garfield’s superior ability to identify spatial niches (Fig. S4c-e). While single-cell multi-omics methods like MultiVI and MOFA+ struggled to distinguish niches, and spatial multi-omics methods like NicheCompass and SpatialGlue showed limited performance in fine tissue niches (e.g., islm), Garfield effectively resolved detailed niche structures, including ctx layers, ccg/aco, ls, acb, and vl. Importantly, Garfield uniquely identified the sophisticated niche structure of islm, which competing algorithms failed to detect. Moreover, our findings indicate that Garfield surpasses single-modal approaches by effectively integrating multi-modal information, thereby enhancing the quality of representation learning (Fig. 2j).

Additionally, Garfield exhibited robust performance across varying neighbor counts and random seeds (Fig. S5). These results underscore Garfield’s potential in cross-modality representation, spatial niche identification, and downstream visualization, establishing it as a powerful tool for spatial multi-omics integration.

### 2.4 Garfield Outperforms State-of-the-Art Tools for Tissue Niche Detection

To evaluate Garfield’s performance in tissue niche identification, we compared it to other spatial representation methods, including NicheCompass[24], CellCharter[22], and GraphST[20], focusing on cellular representations and niche labels. The bench-marking was conducted on a Slide-seqV2 dataset of the mouse hippocampus, known for its well-defined spatial tissue organization that closely aligns with anatomical structures[6].

Since this dataset lacks ground truth labels, we qualitatively compared Garfield’s identified niches to an anatomical map from the Allen Mouse Brain Reference Atlas[41] (Fig. 3a). Garfield’s niches showed strong alignment with anatomical subcomponents, accurately reflecting local tissue structures (Fig. 3b-c). Manually annotated niches displayed unique spatial distributions (Fig. S6a) and were associated with known markers (Fig. S6b), validating their use as pseudo-ground truth for benchmarking. Additionally, niches with similar anatomical structures exhibited comparable variations in cell type abundance (Fig. S6c), suggesting that differences in cell type composition, rather than gene expression alone, contribute to biological variation within niches.

**Fig. 3.**
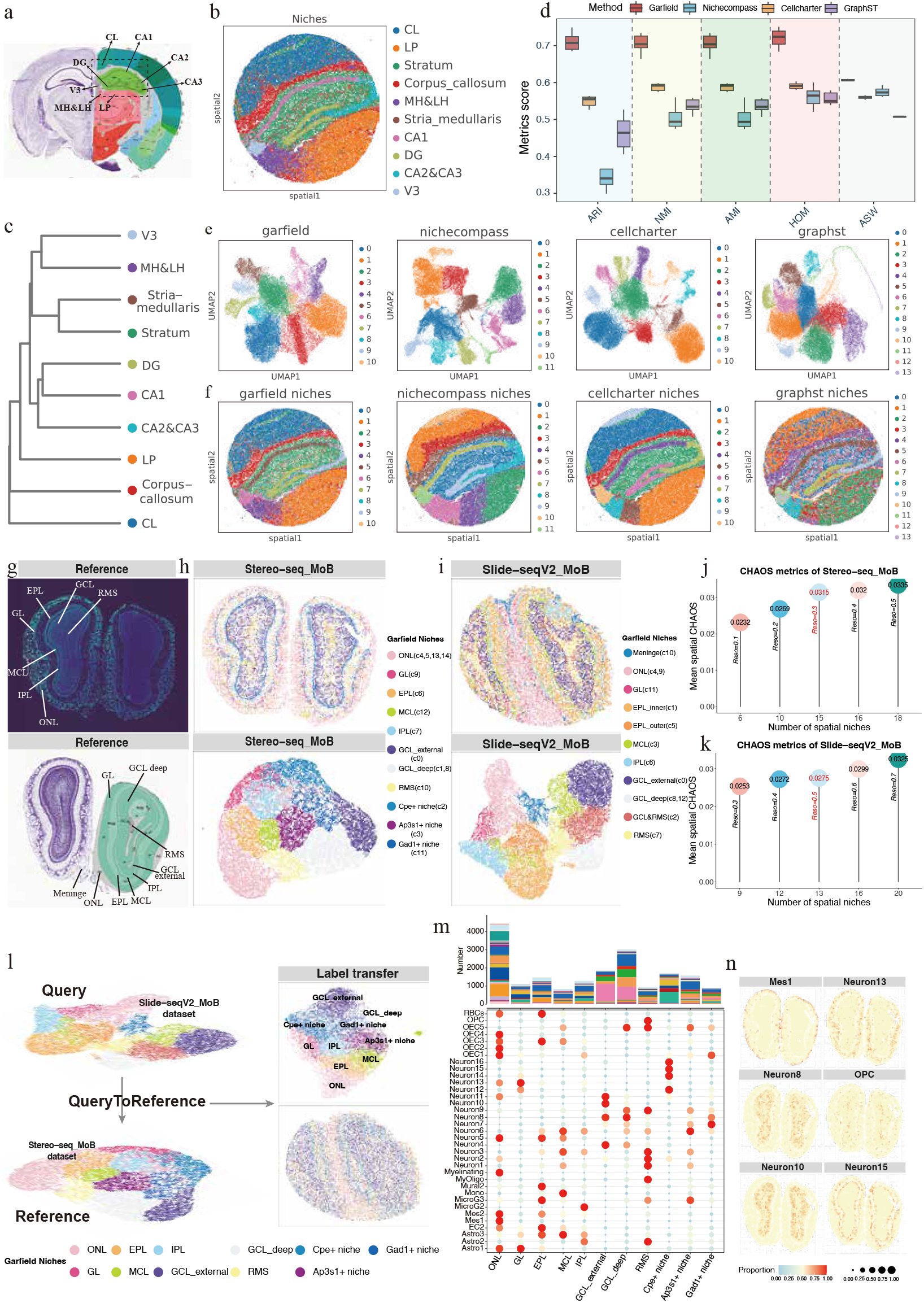
Garfield accurately identifies niches across technologies and tissues. **a**. Anatomical regions include CL (corpus callosum), LP (lateral posterior thalamic nucleus), Stratum, MH&LH (medial and lateral habenula), Stria medullaris, DG (dentate gyrus), CA1, CA2&CA3, and V3, annotated according to histological and spatial organization from Allen Reference Atlas. **b**. Manual annotation for spatial niches with clear boundaries, reflecting the anatomical regions of the hippocampus. Niches are colored based on their spatial organization. **c**. Dendrogram of niche clustering based on cellular and spatial similarity, showing the relationships among the annotated hippocampal regions. **d**. Box plots of evaluation metrics (ARI, NMI, AMI, HOM, and ASW) demonstrate that Garfield outperforms competing methods (NicheCompass, CellCharter, and GraphST) in clustering accuracy, homogeneity, and spatial organization preservation. **e-f**. UMAP (e) and spatial visualization (f) of niches identified at optimal resolution across various algorithms. **g**. Structural overview of the mouse olfactory bulb, with layers annotated based on the Allen Reference Atlas of the Mouse Brain. The olfactory bulb is organized in an out-to-in pattern, comprising meninges, ONL, GL, EPL, MCL, IPL, GCL, and RMS. The surrounding membrane is termed the meninges. **h**. Spatial niches identified by Garfield in mouse olfactory bulb (MoB) slices sequenced with Stereo-seq, visualized both on the tissue section (physical location) and in a UMAP plot. **i**. Similar to panel h, but the data shown here were derived from MoB slices sequenced with Slide-seqV2. **j–k**. Bar plots illustrating CHAOS values for MoB tissue slices sequenced by Stereo-seq (j) and Slide-seqV2 (k), varying the resolution of pre-specified spatial niches. The highlighted resolution (in red) was selected for subsequent niche discovery. **h–i**. Scatter plots displaying the spatial distribution of essential MoB-related marker genes in slices from Stereo-seq (h) and Slide-seqV2 (i). **l**. UMAP representation of the Garfield spatial reference, with query cells mapped through fine-tuning. **m**. A heatmap showing the estimated cell-type proportions for representative cell types across spatial niches detected by Garfield (Stereo-seq slice). The color scale is normalized to a 0–1 range. The top bar plot shows the abundance of different cell types from panel a across the various niches. **n**. A spatial scatter plot illustrating the distribution of estimated cell-type proportions, derived from label transfer by Garfield, for representative cell types across spatial locations.

Using ARI, NMI, AMI, HOM, and ASW as evaluation metrics, we benchmarked Garfield against competing tools. Garfield consistently outperformed all methods across these metrics (Fig. 3d). While all methods could distinguish major structures like CA1, CA2, CA3, and DG, GraphST struggled to resolve finer niche structures such as LP and Stratum. NicheCompass had difficulty identifying the Corpus callosum and exhibited potential over-segmentation, particularly in the LP niche. Both CellCharter and Garfield suggested LP might be homogeneous, but CellCharter’s performance on Stratum was notably weaker than Garfield’s. Garfield delivered the most robust results, accurately characterizing the spatial distribution of all niches in the mouse hippocampus. These findings demonstrate Garfield’s superior ability to detect tissue niches.

### 2.5 Mapping Shared and Unique Spatial Niches in the Mouse Olfactory Bulb Using Stereo-seq and Slide-seqV2

Garfield’s ability to identify niches across different spatial sequencing platforms was demonstrated using two mouse olfactory bulb spatial transcriptome slices profiled by Slide-seqV2[6] and Stereo-seq[7] (Fig. 3g-i). Garfield accurately resolved known tissue structures, including the olfactory nerve layer (ONL), glomerular layer (GL), external plexiform layer (EPL), mitral cell layer (MCL), granule cell layer (GCL), internal plexiform layer (IPL), and rostral migratory stream (RMS), based on DAPI-stained image[42] and Allen Mouse Brain Atlas annotations[41] (Fig. 3g).

Notably, Garfield detected finer niche structures in Slide-seqV2 data, such as subdividing the EPL into EPL_inner and EPL_outer. This difference likely reflects the higher capture resolution of Slide-seqV2, highlighting Garfield’s capacity to discern detailed spatial structures. The identified niches exhibited specific spatial distributions and distinct marker gene expressions (Fig. S7). For example, niche 10 (Slide-seqV2) was enriched with Ptgds[43] (meninge[44]), niche 4, 9 (ONL) with S100a5[45], niche 6 (IPL) with Slc17a7[46], and niche 7 (RMS) with Mbp[47]. Quantitative analysis[46] confirmed these patterns, showing better spatial continuity and smoothness of niches identified by Garfield across platforms, irrespective of resolution (Fig. 3j-k).

To address the transferability of niches between spatial omics datasets, Garfield used Stereo-seq data as a reference and Slide-seqV2 data as a query. Garfield effectively transferred niche labels from the reference to the query, maintaining consistency in the latent space (Fig. 3l). This result suggests Garfield’s potential for annotation migration across datasets, revealing conserved niches between sequencing technologies. Such findings pave the way for constructing comprehensive 3D spatial maps.

Additionally, we explored cell-type enrichment in different niches using publicly available mouse olfactory bulb scRNA-seq data from GSE121891[45]. Cell type annotations mapped to Stereo-seq data revealed unique niche-specific characteristics, including neuronal subtypes across adjacent layers, mesenchymal cell subpopulations within the ONL, and oligodendrocyte precursor cells (OPCs) in the RMS (Fig. 3m-n). These findings further demonstrate Garfield’s utility in uncovering complex spatial and cellular relationships within tissues.

### 2.6 Uncovering Heterogeneity in the NSCLC Tumor Microenvironment Using Garfield

To demonstrate Garfield’s ability to resolve tumor microenvironment heterogeneity in large-scale cancer datasets, we applied it to a human non-small cell lung cancer (NSCLC) dataset consisting of eight tissue sections (702,199 cells) from five donors, with two donors having two or three technical replicates, respectively. A spatial atlas of NSCLC was constructed using all samples except donor 13 (625,663 cells). Garfield identified 16 distinct niches driven by differential cell composition (Fig. 4a, Fig. S8b left) and spatial organization (Fig. S8b right). Several tumor-specific niches (e.g., niches 0, 1, 5, 9, and 13) were identified as donor-specific but shared across technical replicates, indicating the removal of batch effects with Garfield (Fig. 4c, Fig. S8b). Notably, niche 7 was found across donors, particularly donors 6 and 9, and defined as a donor-shared niche. It exhibited a distinct spatial distribution in histological images and was enriched in epithelial and myeloid cells (Fig. 4d-e). Additionally, donor-specific niches were observed, such as niche 1 (specific to donor 6, Fig. 4f) and niche 13 (specific to donor 12, Fig. 4h). Spatial location and neighborhood cell compositions corroborated these findings (Fig. 4g, Fig. 4i). Key core genes driving these niches were identified, including EGFR in donor 6-specific niche 1 (Fig. S8c) and S100A6 in donor 12-specific niche 13 (Fig. S8d). Other niches, such as niche 2, were co-enriched with tumor-related fibroblasts and neutrophils, suggesting their relevance to tumor progression (Fig. S8a-b). Analysis of cell-type proportions across niches revealed that each niche could be annotated by a significantly enriched cell type, showcasing Garfield’s ability to highlight tumor heterogeneity in NSCLC (Fig. 4j). These findings were further supported by preference analysis using Ro/e (Fig. S8e).

**Fig. 4.**
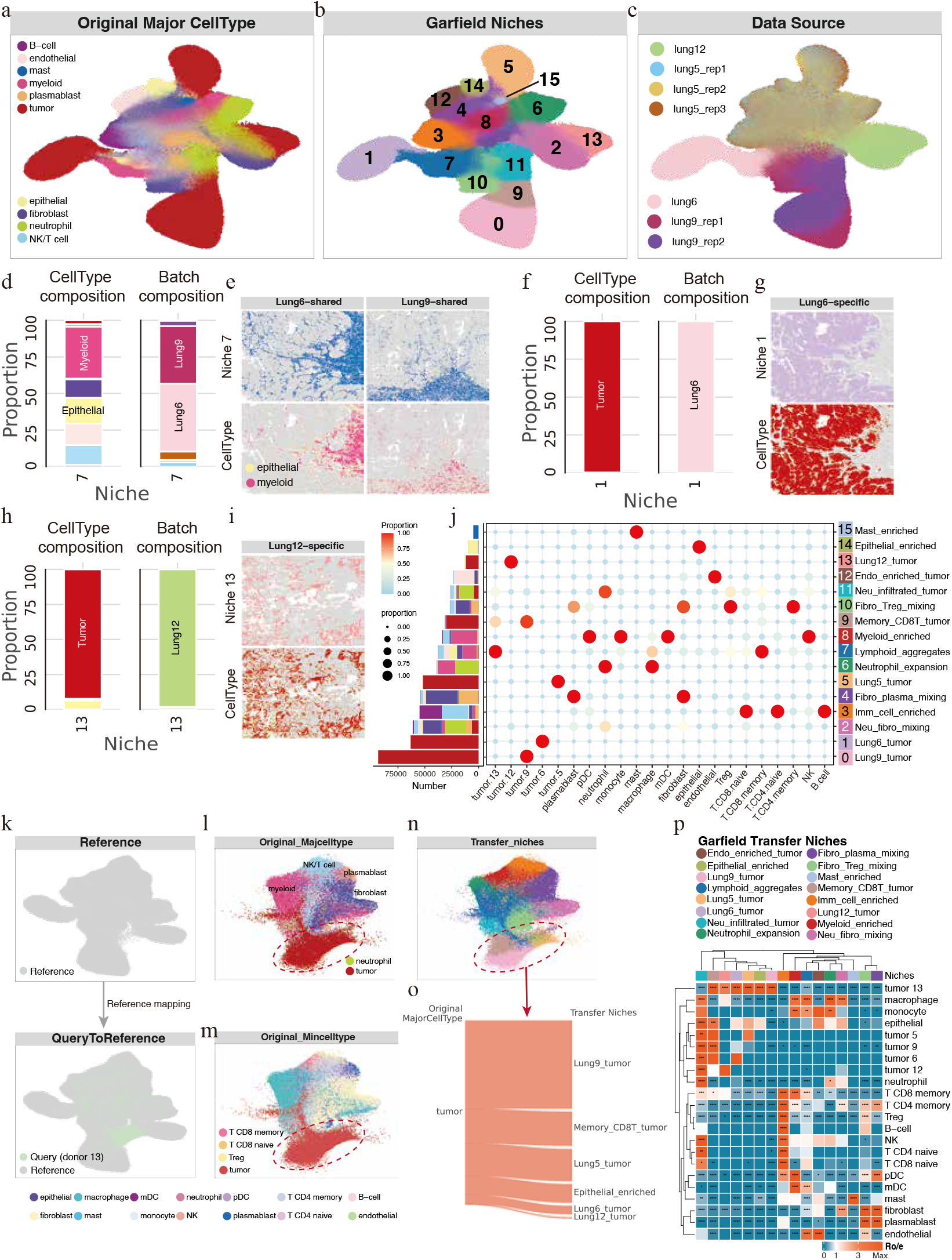
Garfield constructs a spatial atlas of heterogeneous non-small cell lung cancer (NSCLC) patients and characterize changes in district cellular abundance and gene signatures within niches. **a**. UMAP representation of the major cell types, including tumor, fibroblast, myeloid, epithelial, and others. **b**. Same as panel a, UMAP embedding showing Garfield-derived spatial niches. **c**. UMAP visualization highlighting the data sources (patients and replicates). **d**. Proportions of major cell types(left) or data source(right) in donor-shared niche 7. **e**. Spatial distribution of donor-shared niche 7 across heterogeneous samples and the top 2 cell types in corresponding regions. **f**. Proportions of major cell types(left) or data source(right) in donor-specific niche 1. **g**. Tissue sections from Lung 6 highlighting the spatial localization of lung-specific niche and cell type. **h**. Similar to panel f, but from donor-specific niche 13. **i**. Tissue sections from Lung 12 highlighting the specific spatial localization of niche 13 and tumor cell. **j**. Distribution and abundance of cell types across niches, with bubble size representing cell proportion and color indicating niche identity. **k**. UMAP visualization of the Garfield spatial reference(top) and the query dataset(bottom) mapped via transfer learning. **l-n**. UMAPs depicting the original major cell types(l), original minor cell types(m), and the transferred niches(n). **o**. Sankey plot of query data illustrates how the original tumor cell map to the transferred niches. **p**. Preferential localization of spatial transfer niches identified by Garfield, based on the ratio of observed to expected cell numbers (Ro/e). A ratio of 1 is indicated by white, with values >1 signifying enrichment in the specific niche.

To evaluate Garfield’s spatial reference mapping capability, we used data from a new donor (donor 13) and mapped it onto the constructed spatial reference using minimal weight-restricted fine-tuning. A k-nearest neighbors classifier trained on Garfield’s latent space transferred niche labels to the query cells, achieving accurate integration while preserving biological features (Fig. 4k-n, Fig. S8f). Label transfer identified a donor 13-specific niche outside the reference, closely related to a tumor-associated niche, with a consistent cell-type distribution (Fig. 4k-o). This indicates Garfield’s ability to detect novel spatial variations while maintaining correlations between reference and query datasets. Taken together, these results highlight Garfield’s capacity to construct large-scale spatial reference atlas and map query spatial datasets to a single-cell resolution spatial atlas with minimal fine-tuning, enabling the discovery of new spatial patterns in complex datasets.

### 2.7 Garfield Identifies Heterogeneous Niches in Human Breast Cancer 10x Xenium Data

Using Garfield, we analyzed a human breast cancer (BC) dataset profiled with 10x Xenium technology to unravel tumor modular organization and interactions between tumor cells and their microenvironment. Garfield effectively captured biological variation and integrated replicates from the same tissue (Fig. 5a-c). By applying Leiden clustering on Garfield’s latent space, we identified 14 distinct niches with clear boundaries, independent of spatial resolution (Fig. S9a). A closer examination of the niches revealed that each was composed of unique combinations of cell types, underscoring their heterogeneity (Fig. 5a-b, d-e, Fig. S9e). For example, niche 1 and niche 8 were mainly enriched with invasive and proliferating tumor cells, respectively, showing similar spatial distributions across replicates (Fig. 5d-e). We also observed immune cells such as T cells, B cells, and DCs were prevalent in niches 3, 4, and 12, suggesting a close association with tumor development. These niches were further supported by well-known marker gene enrichment, validating the layered structure (Fig. S9b).

**Fig. 5.**
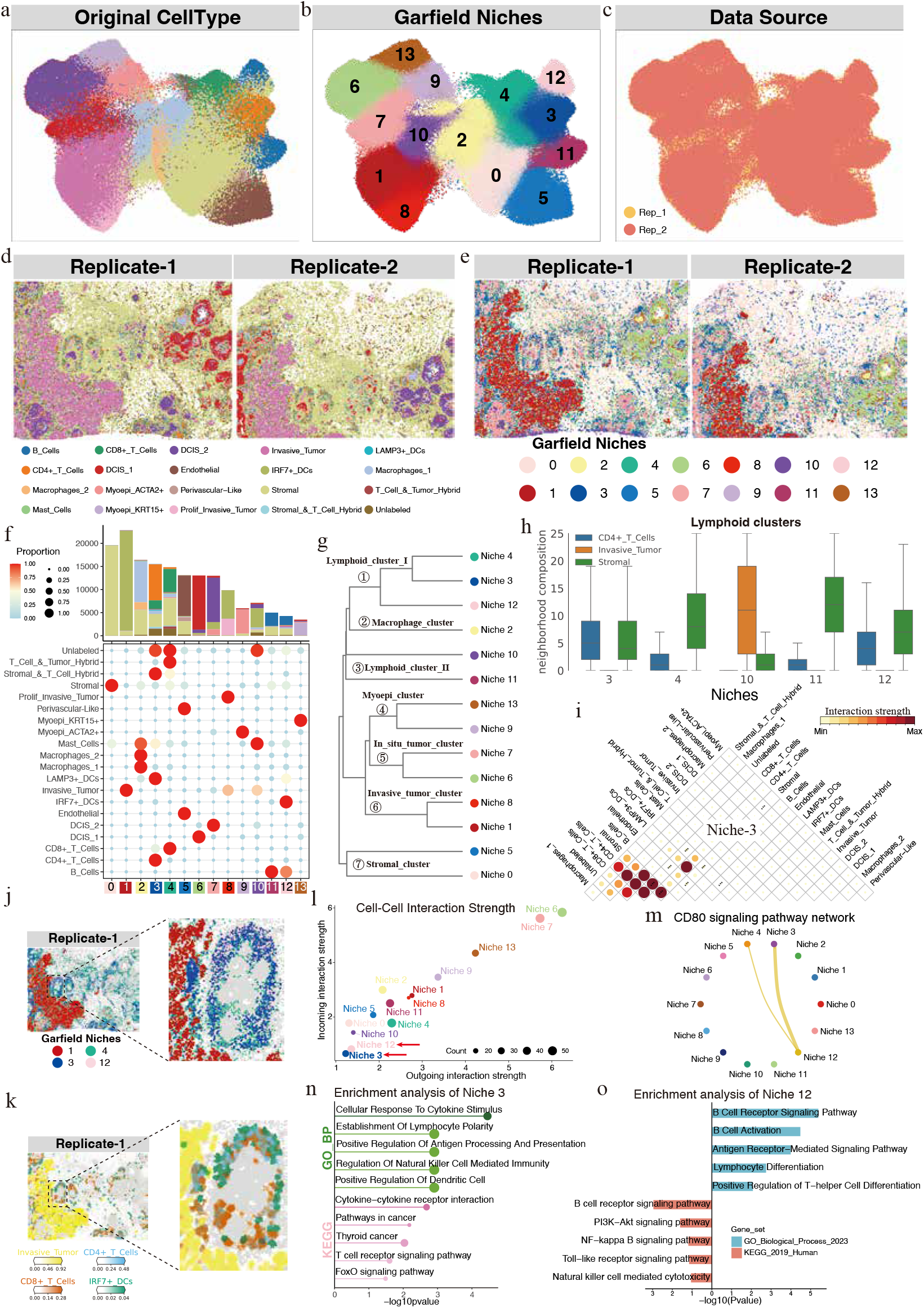
Garfield identifies functionally distinct niches with unique spatial significance. **a**. UMAP representation of the Garfield latent space after integrating two tissue replicates from a 313-probe 10X Xenium BC dataset, colored by the original cell types. **b**. Same as panel a, but colored by the spatial niches identified by Garfield. **c**. Integrated latent space, color-coded by data source, demonstrating the efficacy of the integration approach implemented by Garfield. **d**. Spatial distribution of cells from replicate-1 (left) and replicate-2 (right) of the 10X Xenium BC dataset, with cells colored according to their original types, and spatial locations annotated. **e**. Spatial niches identified by Garfield in both tissue slices. **f**. Cell type distribution and abundance across niches, with bubble size denoting cell proportion and color indicating niche identity. **g**. A dendrogram based on Garfield latent reveals a hierarchy of molecular similar niches. **h**. Neighborhood Composition in Lymphoid-Related Niches. Boxplots depict the distribution of neighboring cell types within lymphoid-related niches, focusing on the 25 closest cells for each niche. The median is marked by the center line, while the box encompasses the interquartile range, with whiskers extending up to 1.5 times this range. To ensure clarity, only cell types contributing an average of 5–60% to the neighborhood composition of any niche are displayed. **i**. Heatmaps showing the proximal cell-cell interactions between cell types within niche 3. The color scale represents the strength of interaction between cell types i and j. Statistical significance was assessed using a permutation test (n = 1,000): * p < 0.05, ** p < 0.01, *** p < 0.001. **j**. Spatial distribution of main tumor-specific niche and lymphoid niches within replicate-1, highlighting their localized enrichment of specific niches. The proximal local network at the tumor edge is magnified and highlighted. **k**. Spatial plot displaying the relative densities of invasive tumor cells (yellow), CD8+ T cells (red), CD4+ T cells (blue) and IRF7+ DCs (green) within the tissue sections, emphasizing their localized enrichment of specific cell types. Gray regions represent background areas. **l**. Cell-Cell interaction strength across niches. Scatterplot displaying the outgoing and incoming interaction strengths of various niches, reflecting the intensity of intercellular communication. **m**. Network visualization of niches involved in the CD80 signaling pathway. Arrows represent signaling interactions, with arrow thickness indicating interaction strength. **n-o**. Functional enrichment analysis highlights the molecular pathways and biological processes associated with Niche 3 (n) and Niche 12 (o).

We further explore cell composition within niches and perform hierarchical clustering based on Garfield latent (Fig. 5f-g). Hierarchical clustering grouped niches with similar cell-type compositions and gene features into different compartments, such as lymphoid_cluster_I (niches 3, 4, and 12), lymphoid_cluster_II (niches 10 and 11), Invasive_tumor_cluster (niches 1 and 8), and so on (Fig. 5g). For lymphoid clusters, we performed neighborhood composition analysis, and confirmed consistent cell types co-localization within lymphoid clusters (Fig. 5i), aligning with previous observations (Fig. 5a-b). To further explore the functional dynamics of lymphoid niches, we constructed proximal interaction networks for niches 3, 4, and 12 (Method). Strong interactions between CD4+ and CD8+ T cells and stromal cells were identified in these three niches (Fig. 5i, Fig. S9c-d), while niche 12 also showed additional crosstalk between B cells and immune/stromal cells, likely due to its high B cell enrichment (Fig. S9d). Given prior evidence that myoepithelial cells can form barriers to inhibit tumor invasion[48], we hypothesize that these lymphoid niches may form a local immune regulatory network to curb tumor progression.

To substantiate this hypothesis, we firstly visualized tumor-specific niche 1 and lymphoid-associated niches 3, 4, and 12 on tissue sections (Fig. 5j). Notably, these lymphoid-related niches demonstrated a consistent enrichment at the periphery of the invasive tumor niche, forming a discernible aggregation network. This spatial pattern was further corroborated through independent replicates (Fig. S9f). At the cell type level, significant enrichment of immune cell populations, including CD4+, CD8+ T, and dendritic cells (DCs), was observed at the same invasive tumor margins, consistent with prior findings (Fig. 5k, Fig. S9g). These results provide preliminary evidence for the co-localization of diverse immune cell types at the tumor boundaries, likely facilitating the formation of a localized immune regulatory network through niches interactions. To further elucidate this phenomenon, we employed CellChat V2[49], an advanced tool for inferring cell-cell interactions with integrated spatial data. Analysis across all niches revealed that niches 3 and 12 exhibited minimal incoming and outgoing interaction strengths, indicative of the establishment of a localized interaction network within these regions (Fig. 5l). Moreover, the CD80 signaling pathway was identified as a key regulatory axis driving this local network, underscoring its potential role in modulating immune-tumor dynamics (Fig. 5m). To investigate the underlying mechanisms of this local network at a functional level, we conducted enrichment analysis on the differential markers of niches 3 and 12 (Fig. 5n-o). Both niches were found to be enriched in tumor-related pathways, including antigen-mediated signaling pathway and cell-mediated immunity. Additionally, distinct signaling pathways were identified for each niche, highlighting their unique characteristics. For instance, niche 3 was specifically associated with the cellular response to cytokine stimulus (Fig. 5n), while niche 12 was linked to B cell activation and related processes (Fig. 5o). These findings indicate that while niches 3 and 12 share core functional features, contributing to network synergy, they also possess niche-specific attributes. This balance between shared and unique functions is essential for maintaining the functional diversity and adaptability of the network, thereby supporting its scalability and effectiveness in regulating immune-tumor interactions.

## 3 Discussion

Our findings demonstrate that Garfield, a novel and versatile single-cell and spatial embedding algorithm, provides a robust framework for integrating uni- and multi-modal single-cell and spatial omics data. By leveraging variational graph autoencoder and a self-supervised, multi-task learning approach, Garfield successfully unifies molecular and spatial information, enabling the precise characterization of cellular landscapes and tissue-specific niches. This capability marks a significant advancement over existing methods, such as Seurat V4, MultiVI, and MOFA+, which rely on more limited (graph) structures and often struggle with spatial and cross-modality integration. The benchmarking results highlight Garfield’s scientific implications, as it consistently outperformed state-of-the-art tools across various datasets, including scRNA + scATAC, scRNA + scADT, and spatial multi-omics.

Particularly in spatial niche detection, Garfield excelled in identifying fine-grained anatomical and functional structures, such as the olfactory bulb’s internal plexiform layer, tumor-specific niches in NSCLC datasets and proximal cell-cell interaction network. Garfield revealed novel insight on breast tumor niches by identifing immune niches at tumor margins orchestrating localized CD80-mediated interactions between T cells, B cells, and dendritic cells. These niches formed barrier-like structures around invasive tumor regions, with lymphoid niches enriched for cytokine response (e.g., IFN-) and B cell activation pathways. Spatial interaction analysis revealed conserved CD80 signaling hubs across replicates, suggesting a coordinated immune surveillance mechanism. These findings underscore its utility in uncovering heterogeneity within complex tissue environments and its ability to integrate information across modalities and resolutions, setting the stage for broad applications in biomedical research.

Garfield’s ability to map query datasets to reference atlases without retraining positions it as a platform for collaborative spatial atlas initiatives. Its capability to construct large-scale spatial reference atlases and seamlessly map query datasets enables a deeper understanding of conserved spatial niches across platforms and conditions. By enabling joint analysis of single-cell and spatial data, Garfield opens avenues to explore conserved architectural principles across tissues and diseases. Extensions to temporal multi-omics or 3D spatial mapping could further illuminate dynamic cellular ecosystems.

Despite these achievements, some limitations remain. While Garfield efficiently integrates multi-modal data and resolves batch effects, its performance in highly sparse datasets and extreme high-dimensional spaces requires further optimization. Additionally, the reliance on reference atlases, such as the Allen Brain Atlas, for validating niche detection introduces potential biases when analyzing previously uncharacterized tissues. Future efforts could focus on enhancing Garfield’s ability to function independently of existing references while improving computational efficiency for large-scale datasets.

Looking ahead, advancing the field of spatial and single-cell multi-omics analysis will require addressing key challenges such as improved scalability, universal applicability across technologies, and deeper insights into dynamic cellular processes. In summary, Garfield bridges a long-standing divide between single-cell and spatial omics, offering a unified framework for hypothesis generation and validation. By providing deeper insights into cellular heterogeneity and spatial organization, it holds immense potential for advancing the understanding of complex biological systems and developing next-generation applications in precision medicine and tissue engineering.

## 5 Methods

### 5.1 Modeling Architecture

#### 5.1.1 Data Loading and Preprocessing

The input to the Garfield model can include one or multiple datasets from various omics sources (such as single-cell RNA, ATAC, ADT, or spatial data), where each row corresponds to a cell and each column represents a feature. Following data loading, Garfield processes the dataset by removing non-informative genes, thereby optimizing the training process. Firstly, genes expressed in fewer than three cells and cells expressing fewer than 100 features are excluded by default. Then, raw counts undergo library-size normalization followed by log transformation. In the case of ATAC and ADT data, term frequency-inverse document frequency (TF-IDF) and Centered Log-Ratio (CLR) normalization methods are applied.

Subsequently, the top 3,000 highly variable genes (HVGs) are selected by default and used for dimensionality reduction via PCA. For chromatin peak data, the top 10,000 highly variable peaks (HVPs) are selected by default, with latent semantic indexing (LSI) employed for dimensionality reduction. Notably, in multi-omics datasets, features derived from PCA or LSI are used as node attributes within the encoder model. This approach has demonstrated exceptional performance in representation learning, as highlighted in the previous study [50]. While in the unimodal scenario, the default is to use original expression profiles as the input of the model.

#### 5.1.2 General dataset

A general `“GraphAnnTorchDataset”` class, inspired by NicheCompass [24], is implemented in Garfield to process single-cell or spatial omics data, enabling graph-based learning by extracting node features, adjacency matrices, and batch labels. In this framework, let 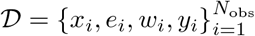 represents the dataset, where *N*_obs_ is the total number of observations. Here, 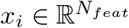 denotes the specific omics feature vector for observation *i*, with *N* features *N*_*feat*_, and 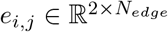 represents the connectivity between observation *i* and its neighbor observation *j*, as derived from the adjacency matrix **A**. The edge weight *w*_*i,j*_ indicates the connectivity weight between observation *i* and its neighbor observation *j*, which is also extracted from **A**. The *y*_*i*_ ∈ ℝ represents the batch label for observation *i*. The features 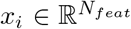 represents the expression profile of observation *i* obtained from `“adata.X”` or `adata.obsm[“feat”]`, and can consist of either raw counts or processed features, supporting the integration of multiple data types (e.g., unimodal or multimodal scenarios).

In the unimodal scenario, we define 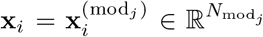,where 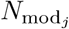 is the number of features from the specific *j* modality, and *j* refers to one of three common modalities (RNA, ATAC, or ADT). In the multimodal scenario, **x**_*i*_ is a concatenated vector of processed gene expression and chromatin accessibility or protein features.

For chromatin peaks, this is represented as 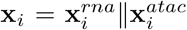 with 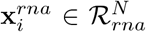,and 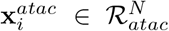, where *N*_*rna*_ and *N*_*atac*_ is the number of genes and peak regions, respectively. For protein features, it is expressed as 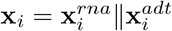 with 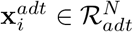, where *N*_*adt*_ is the number of antibody. The adjacency matrix 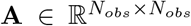 is derived from `“adata.obsp[“connectivities”]”.`

#### 5.1.3 Construction of neighbor graph

Consider a neighborhood graph 𝒢= {*v*_*i*_, *s*_*i*_, *x*_*i*_, *γ*_*i*_}, where each node *v*_*i*_ represents an observation, with the adjacency matrix *s*_*i*_ encoding its connectivity to single-cell or spatial neighbors. From this matrix, we can derive the edge relationships and their associated weights between *v*_*i*_ and other observations. The vector *x*_*i*_ denotes the expression vector of node *v*_*i*_, while *γ*_*i*_ represents its label vector. In the Garfield algorithm, 𝒢 is an un-directed neighborhood graph, composed of sample-specific symmetric k-nearest neighbor sub-graphs 𝒢_*k*_, where *k* is the number of samples. The construction of 𝒢depends on the data type.

For single-cell unimodal data, 𝒢 is computed using the `“sc.pp.neighbors”` function from the Scanpy [27] package. In the single-cell multimodal scenario, 𝒢 consists of intra-modality connectivity (intra_connect) and inter-modality connectivity (inter_connect). The calculation of intra_connect is similar to that of the single-cell unimodal case. Inspired by previous studies [51], the connectivity between different modalities are modeled as a linear assignment optimization problem, and inter_connect is obtained by solving this objective function. We extended the original cell-matching method [51, 52] to support not only scRNA-seq and single-cell proteomics but also scRNA-seq and scATAC-seq data. Our approach involves scoring the gene activity in scATAC-seq data prior to initial matching via linear assignment, ensuring that peak features overlap with those of scRNA-seq data. This step is automatically performed by Garfield’s built-in “gene_scores” function, which was modified by SIMBA [53]. Thus, the final connectivity are computed as the weighted sum of the intra_connect (ℰ_*intra*_) and inter_connect (ℰ_*inter*_). Mathematically, this can be expressed as:

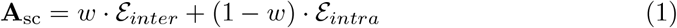

where the weight *w* represents the relative contributions of ℰ_*inter*_ and ℰ_*intra*_ to the overall connectivity.

For spatial single- or multi-modal data, Garfield provides four methods to compute spatial connectivity (E_spatial_): mu_std, radius, KNN, and Squidpy.

1. mu_std: refer to previous study [23], Garfield computes a full pairwise Euclidean distance matrix between cells based on their spatial coordinates. For each cell, the k-nearest neighbors are identified, and their corresponding distances are analyzed. The boundary for establishing a connectivity is defined as boundary = *µ*+*σ*, where *µ* represents the mean and *σ* the standard deviation of the distances to the nearest neighbors. If the distance between two cells is within this boundary, an edge is assigned a weight of 1; otherwise, the weight is 0. This is formalized as:

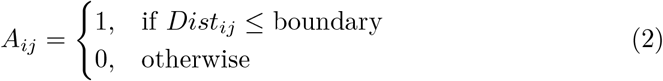
2. Radius: in the Radius method, a radius-based neighbor search is employed. A `“NearestNeighbors”` object is instantiated with a specified radius parameter, and the function identifies all neighbors for each cell within this radius. The resulting cell pairs are connected in the graph, with edge weights corresponding to their spatial distances.
3. KNN: in the KNN method, the k-nearest neighbors of each cell are identified using the `“kneighbors”` function, which returns both the distances and the indices of the nearest neighbors. Graph edges are then established between each cell and its k-nearest neighbors, with edge weights representing the spatial distances.
4. Squidpy [28]: Garfield offers an additional method implemented through Squidpy’s `“spatial_neighbors”` function, which computes spatial connectivity based on physical proximity.

Additionally, based on the spatial single-modal expression profile, we also compute molecular connectivity (ℰ_molecular_) using the “sc.pp.neighbors” function similar to single-cell data. This function builds a neighbor graph based on expression data, where each node represents a cell, and edges are determined by similarity in molecular profiles. For spatial multi-modal data, we build upon the aforementioned workflow by adapting established single-cell multi-omics processing methods. Utilizing our optimized cell matching algorithm, we identify cross-modality connectivity within spatial multi-modal datasets. These resulting molecular connectivity graph captures both intra-modality (ℰ _*intra*_) and inter-modality (ℰ _*inter*_) relationships (if spatial multimodal data exists), providing a detailed view of interactions at the molecular and spatial levels. As a result, the overall connectivity is computed as the sum of the weights from both spatial connectivity and molecular connectivity. Mathematically, the adjacency matrix **A**_*sp*_ for spatial uni-modal data is defined as:

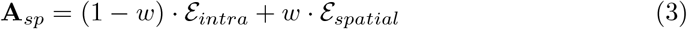

For spatial multi-modal data, it is expressed as:

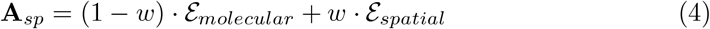

where ℰ_*molecular*_ is defined by:

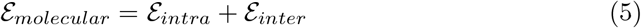

### 5.2 Garfield Model

#### 5.2.1 Model overview

To address the scarcity and task-specific nature of label information, we propose learning embeddings in an unsupervised manner to enhance model generalizability. Garfield begins by encoding all cells into a unified graph, where nodes represent individual cells and edges capture inter-cellular relationships. It then applies a powerful representation learning approach, the Variational Graph AutoEncoder (VGAE) [25], incorporating a SVD-based graph contrastive strategy, an optional MMD-based transformation, and a multi-task learning setup. This setup includes both node- and edge-level tasks, embedding cells into a low-dimensional, batch-invariant biological space ideal for batch effect correction and multi-omics integration at the single-cell and spatial levels.

The model is built on an encoder-decoder framework. Specifically, the encoder consists of a graph encoder module, while the decoder includes a graph decoder module to reconstruct the adjacency matrix from the latent vector, and a feature reconstruction decoder module for handling omics modalities. This dual-module structure ensures that the latent representations capture both physical graph and molecular information while integrating spatial data.

Inspired by the LightGCL [33] framework, Garfield uses a GAT backbone to automatically extract local graph dependencies. The SVD-guided augmentation enhances graph contrastive learning by analyzing global collaborative relations, improving single-cell representation. Additionally, an optional drop edge augmentation strategy is available.

In line with variational autoencoder standards, we use a standard normal distribution as the prior for latent random variables, denoted as *Z*∼𝒩 (0, 1), and apply the reparameterization trick, enabling end-to-end training through backpropagation.

#### 5.2.2 Data augmentation

Contrastive learning aims to capture invariant representations between similar and dissimilar data pairs. To create similar pairs, we apply data augmentation to generate perturbed views of the input data. We offer two graph augmentation methods: ApproxSVD-guided augmentation [33] (default) and edge-drop augmentation [54].

For ApproxSVD-guided augmentation, we perform a sparse approximation of Singular Value Decomposition (ApproxSVD) on the original adjacency matrix *A* as *A* = *USV* ^⊤^. Here, *U* and *V* are orthonormal matrices representing the eigenvectors of the row-row and column-column correlation matrices of *A*, while *S* is a diagonal matrix of singular values. The largest singular values typically correspond to the principal components of the graph. We truncate the singular values, keeping only the top *q*, and reconstruct the adjacency matrix as 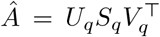,where *U*_*q*_ ∈ *R*^*m*×*q*^ and *V*_*q*_ ∈ *R*^*n*×*q*^ contain the first *q* columns of *U* and *V*, respectively, and *S*_*q*_ ∈ *R*^*q*×*q*^ is the diagonal matrix of the top *q* singular values. The reconstructed matrix *Â* is a low-rank approximation of *A*, where rank(*Â*) = *q*. The benefits of SVD-based augmentation are twofold: it emphasizes principal components, highlighting critical cell-cell interactions, while preserving global collaborative signals across cell pairs. To handle large adjacency matrices efficiently, we use the randomized SVD algorithm from Halko et al. study [55], which approximates the input matrix using a low-rank orthonormal matrix and then applies SVD on the smaller matrix:

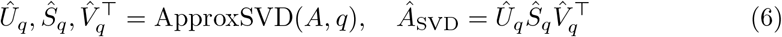

where *q* is the desired rank, and *Û*_*q*_ ∈ ℝ^*m*×*q*^, *Ŝ*_*q*_ ∈ ℝ^*q*×*q*^, and 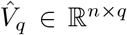 are the approximated matrices.

For edge-drop augmentation (optional), we randomly remove edges from ℰ based on a dropout ratio *p*_drop_. Specifically, we sample an indicator matrix *R*_edge_mask_ ∈ ℝ^*N* ×*N*^ to determine which edges to mask. Each entry *R*_*ij*_ is sampled as:

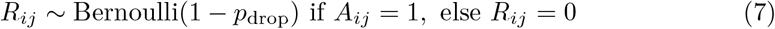

The augmented adjacency matrix is then computed as:

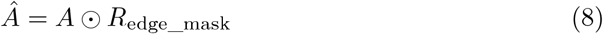

where ⊙ denotes the Hadamard product.

Both ApproxSVD-guided and edge-drop augmentations generate perturbed adjacency matrices *Â* that retain the structure of the original graph. To further enhance robustness, we introduce a feature masking vector *R*_feat_mask_ ∈ ℝ^*N* ×1^ (denoted as *R*_fm_), where each entry *R*_*i*_ ∼ Bernoulli(1− *p*_drop_), with *p*_drop_ as the masking probability. The masked feature matrix is expressed as:

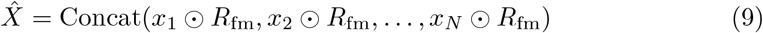

#### 5.2.3 Garfield Encoder

Through data processing and ApproxSVD-guided or edge-drop augmentation strategies, Garfield could generate an original view 𝒢_1_ as well as augmented view 𝒢_2_, we then learn cell embeddings using node-level graph contrastive learning. Given that GAT [31] is a state-of-the-art architecture for graph representation learning and excels at extracting rich embeddings from graph-structured data, we employ it as the feature extractor to capture the latent embedding of the cell nodes. Specifically, in GAT, each node updates its representation by attending to its neighbors, using its own representation as the query. This approach extends the standard neighbor averaging or max-pooling by allowing nodes to compute a weighted average, prioritizing relevant neighbors [56].

For a set of node features *h* = [*h*_1_; *h*_2_; … ; *h*_*N*_], where *h*_*i*_ ∈ ℝ^*D*^, a shared linear transformation with a weight matrix *W* is applied to all nodes. An attention mechanism *α* is then applied pairwise to the transformed node features via *e*_*ij*_ = *α*(*Wh*_*i*_, *Wh*_*j*_), which reflects how much node *j* contributes to node *i*. The attention coefficients for head 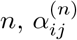,are computed as:

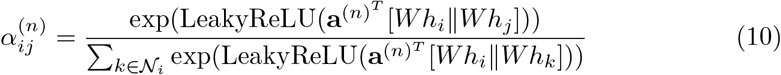

where ∥ denotes concatenation and LeakyReLU is used to allow a small gradient when the argument is non-positive. The normalized coefficients are then combined with the neighboring features (often after applying a non-linearity *σ*) to produce the output node features.

Multi-head attention, denoted as *K*, can be applied in two ways:

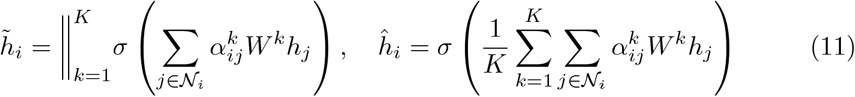

The left formulation is suitable for hidden layers, as it outputs concatenated representations of dimension *K · D*, while the right formulation, which averages the attention heads, is appropriate for the output layer.

In addition to GAT, we also provide GATv2 [30], an improved version with a dynamic attention mechanism. The attention coefficients in GATv2 are calculated as:

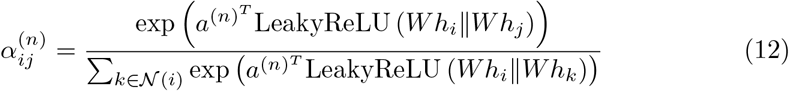

For each node *i*, the updated means in layer *l* are given by:

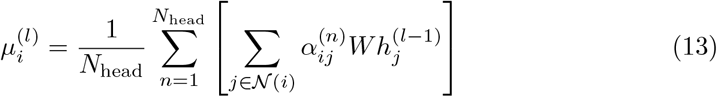

where 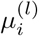 and 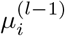 represent the means in the *l*-th and (*l* −1)-th layers, respectively, and 𝒩 (*i*) refers to the neighbors of node *i*. Stacking *L* graph layers results in *L* outputs [*µ*_1_, *µ*_2_, …, *µ*_*L*_], from which we compute the final means and variances as:

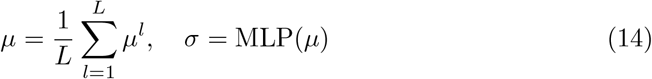

The variances are learned via a one-layer MLP: *σ* = exp(*µ***W** + **b**), where **W** ∈ ℝ^*d*×*d*^ and **b** ∈ ℝ^*d*^ are learnable parameters. To generate latent representations *z*_*i*_, we sample from 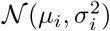 using the reparameterization trick: *z*_*i*_ = *µ*_*i*_ + *σ*_*i*_ ·*ε* where *ε* ∼ *𝒩* (0, **I**) is Gaussian noise.

For Garfield Light, a low-memory version of the model, we replace the graph attention layers with graph convolutional layers [32] for message passing. Additionally, to reduce the dimensionality of *x*_*i*_, an optional fully connected layer with hidden size *N*_hid_ can be added before the graph encoder network to lower memory consumption. Its node-wise formulation is:

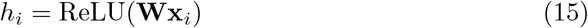

where 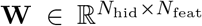 is a learnable weight matrix (bias terms are omitted for simplicity).

#### 5.2.4 Garfield Decoder

After estimating the probability distribution of the latent variables *Z*, the goal of graph generation is to reconstruct both the original graph structure and expression profiles. To achieve this, the Garfield decoder consists of a graph decoder module and an omics decoder module. The omics decoder module is further divided into two sub-modules: the inner product decoder and the feature reconstruction decoder.

For the graph decoder module, the reconstructed adjacency matrix *Ã* is computed using the cosine similarity between pairwise latent features, formulated as:

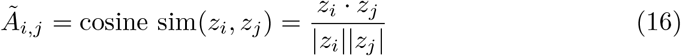

In the omics decoder module, we firstly use the inner product decoder to reconstruct the graph’s adjacency matrix. Given the node embeddings *Z* ∈ ℝ^*N* ×*d*^, where *N* is the number of nodes and *d* is the embedding dimension, the inner product decoder predicts edges by calculating the dot product between node embeddings. The predicted adjacency matrix *Ã*^′^ is expressed as:

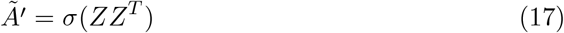

where *Z* ∈ ℝ^*N* ×*d*^ is the embedding matrix, with each row representing a node’s embedding. The product *ZZ*^*T*^ results in an *N* × *N* matrix, where each element 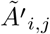 represents the predicted probability of an edge between nodes *i* and *j*. The activation function *σ* (commonly sigmoid) constrains output values to the range [0, 1], making them interpretable as edge probabilities.

The second sub-module, the feature reconstruction decoder, aims to reconstruct the omics expression matrix 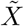 from the embeddings *Z*. Given two nodes *i* and *j* with embeddings *h*_*i*_ and *h*_*j*_, the attention coefficient between them is computed as:

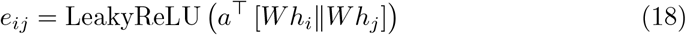

where *a* is a learnable attention vector, *W* is the weight matrix, and ∥ denotes concatenation. The normalized attention coefficients are then calculated using the softmax function:

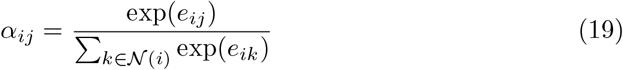

The node embedding is updated as a weighted sum of its neighbors’ embeddings:

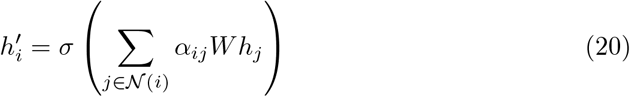

where *σ* is an activation function (e.g., ReLU), and 𝒩 (*i*) is the set of neighboring nodes of *i*. If edge weights *w* are used, they scale the node features during message passing:

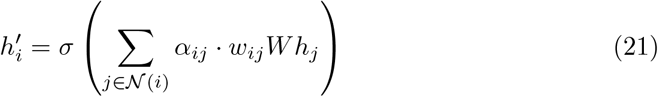

where *w*_*ij*_ is the edge weight between nodes *i* and *j*. By stacking multiple GAT layers and applying attention-based message passing, the decoder reconstructs node features from the latent representations learned during encoding.

#### 5.2.5 Garfield Data Loaders

To scale Garfield to large datasets, where neighborhood graphs 𝒢 can become computationally intensive, we use mini-batch training with inductive neighbor sampling [56]. Unlike methods that load the entire graph into memory [20, 57, 58], which is infeasible for large datasets, we sample a fixed number of neighbors *n*_sam_ = 3 from the *k*-nearest neighbors in 𝒢 for message passing during training.

Similar to NicheCompass [24], we employ two separate task-specific data loaders: a node-level loader for node-level tasks and an edge-level loader for edge-level tasks. Each model iteration includes a forward pass through each loader and a joint backward pass for simultaneous gradient computation. The node-level loader is responsible for omics prediction tasks, such as reconstructing 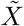 and the adjacency matrix *Ã*. The edge-level loader is used solely for reconstructing *Ã*.

#### 2.5.6 Loss function

The Garfield loss function is composed of edge-level and node-level losses because of edge-level and node-level tasks. The edge-level loss utilizes a weighted binary cross-entropy (BCE) to reconstruct edges in the adjacency matrix *Ã*, while the node-level loss consists of several components: the mean squared error (MSE) for feature reconstruction 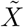,contrastive loss for latent vectors, the Kullback-Leibler divergence (KL) between the variational posteriors and standard normal priors of latent variables, and the batch effect alignment loss for omics data (when multiple batches are present). Furthermore, to ensure the model captures edge-related information in the node-level tasks, a BCE loss with negative sampling is also applied. Among these, the graph reconstruction loss (BCE) and KL divergence loss are jointly referred to as the Evidence Lower Bound (ELBO).

For edge-level loss, Garfield firstly calculates edge reconstruction using a weighted BCE approach to ensure numerical stability. The positive weight 𝒲_pos_ is determined by the ratio of negative to positive edges:

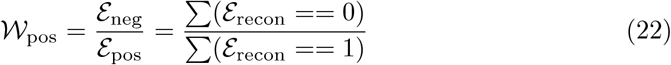

This weighting prevents bias toward negative samples. The mini-batch binary cross-entropy loss for edge reconstruction is given by:

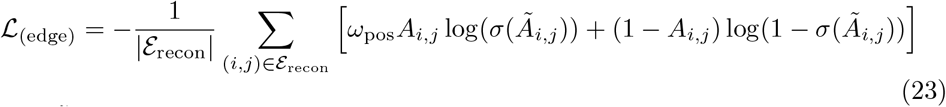

where *Ã* is the edge reconstruction logits matrix, and 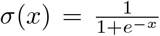 is the sigmoid function applied to logits.

Garfield learns the reconstructed expression profile using a self-supervised loss that combines the mean square error (MSE) from the latent embedding (*Z*) with the omics loss function:

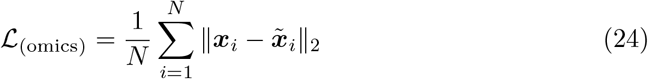

where ***x***_*i*_ and 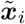 represent the original and reconstructed expression profiles, respectively, *N* is the total number of cells, and ∥ · ∥ _2_ denotes the L2 norm (Euclidean distance), which measures the magnitude of the difference between the original and reconstructed profiles.

For contrastive learning [59, 60] of node-level task, latent embeddings *Z*_1_ (from the original view) and *Z*_2_ (from the augmented view) are projected into a common latent space using a weight-sharing projection head *g*_*I*_ (·), implemented as a two-layer multilayer perceptron (MLP). Given a mini-batch of size *N*, Garfield generates 2N samples 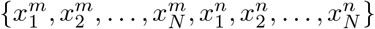 through augmentations (such as SVD-based or edge-drop strategies), forming pairs of original and augmented samples 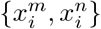.

The similarity between pairs is computed using cosine distance:

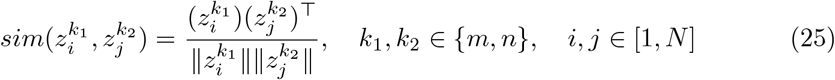

The contrastive loss for a sample 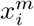 is given by:

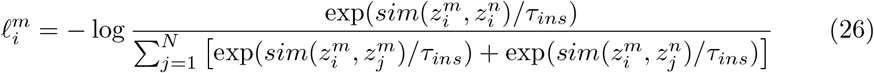

where *τ*_*ins*_ is the temperature parameter for instance-level similarity. The overall instance-level contrastive loss [61] is computed as:

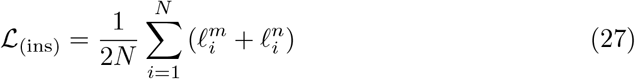

By minimizing this loss, positive pairs are drawn closer while negative pairs are pushed apart, preserving biological variability and implicitly correcting batch effects [58].

In addition to instance-level loss, Garfield also employs cluster-level contrastive loss [62], which has shown robust performance in representation learning. Using a two-layer MLP *g*_*C*_(·), the feature matrix is projected into an *M* -dimensional space (where *M* is the number of clusters). The similarity between cluster pairs is also measured using cosine distance:

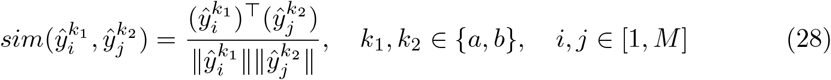

The cluster-level loss for a given cluster 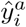 is formulated as:

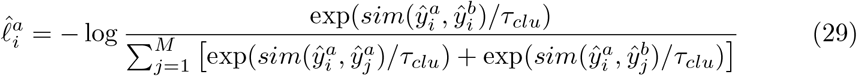

where *τ*_*clu*_ is the temperature parameter for cluster-level similarity. The overall cluster-level contrastive loss is:

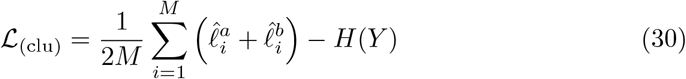

with entropy *H*(*Y*) of cluster assignment probabilities defined as:

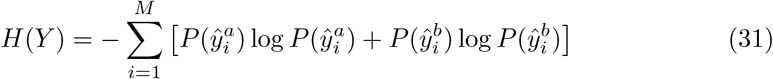

and the probability 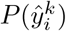 for a mini-batch under each augmentation calculated by:

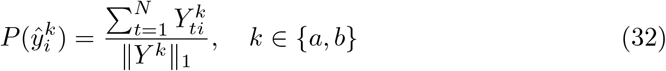

where *Y* ^*a*^ ∈ ℝ^*N* ×*M*^ represent the output of *g*_*C*_(·) for a mini-batch under the original view, with *Y* ^*b*^ denoting the corresponding output under the augmented view. This term *H*(*Y*) helps avoid trivial solutions, ensuring a balanced cluster assignment [63]. The mini-batch KL divergence [24] is composed of two parts: the node-level KL divergence, which involves nodes from the node batch, and the edge-level KL divergence, which includes nodes from both positive and negative edge pairs in the edge batch:

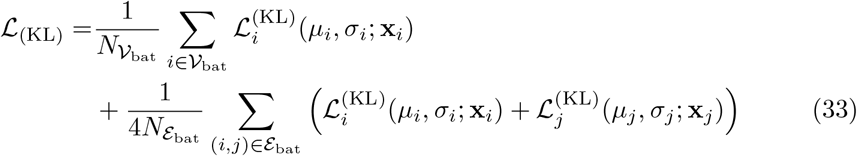

Here, the variational parameters *µ*_*i*_ represents the estimated mean of the approximate posterior normal distribution, obtained as outputs from the graph encoder module, and *σ*_*i*_ represents the estimated standard deviation through one-layer MLP projection based on 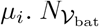 represents the number of nodes in the node batch 𝒱_bat_, while 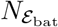 denotes the number of edges in the edge batch ℰ_bat_.

Additionally, a similar edge reconstruction loss 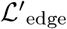 is applied in the node-level tasks using BCE with negative sampling, where *ŷ*^pos^ represents the decoder output for positive (real) edges, *ŷ*^neg^ for negative (randomly sampled) edges, and ℰ is the total number of edges:

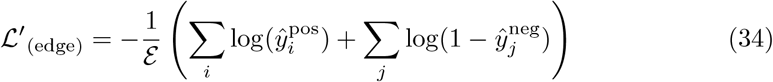

If the input data contains multiple batches, Garfield employs the maximum mean discrepancy (MMD) [34] loss to align data across different batches. Inspired by scArches [26], Garfield uses MMD to match latent distributions *q*_*ϕ*_(*Z*| *s* = *i*) and *q*_*ϕ*_(*Z* |*s* = *j*), where *s* represents the batch key. The MMD loss is formulated as follows:

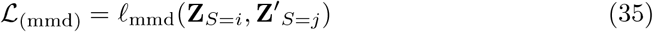

where

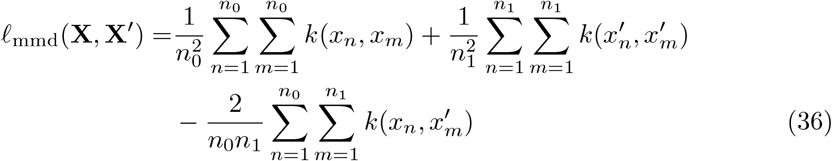

Here, **X** and **X**^′^ are high-dimensional observations under conditions *S*_*i*_ and *S*_*j*_, respectively.

The overall loss function of the Garfield model that is optimized during model training is thus:

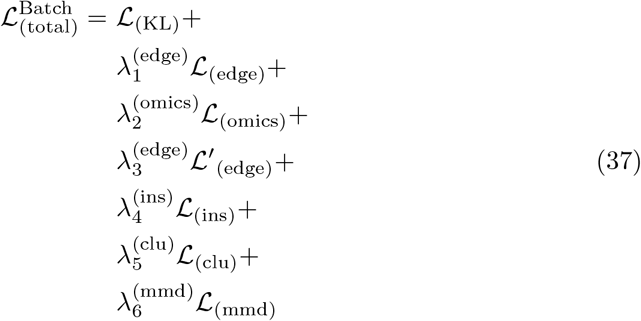

If no batch information is present, the total loss as follows:

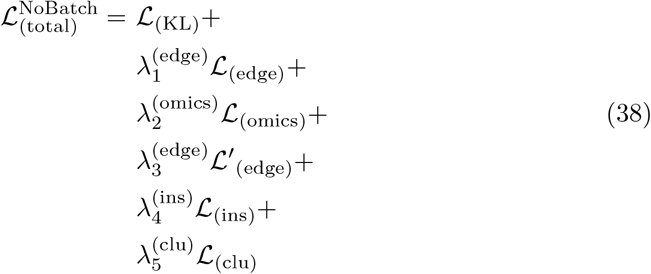

where the weighting factors for the total loss components are denoted as *λ*.

### 5.3 Benchmarking

All benchmarking experiments are fully reproducible and can be accessed through the notebooks provided at https://github.com/zhou-1314/Garfield-reproducibility. These experiments were conducted on a high-performance computing infrastructure equipped with an NVIDIA A800-SXM4-80GB GPU, ensuring robust and efficient execution.

#### 5.3.1 Datasets overview

The datasets presented in this manuscript can be broadly categorized into two groups: single-cell multi-omics datasets and spatial omics datasets. Unless otherwise specified, cell type annotations and metadata were sourced directly from the original publications.

##### Single-cell multi-omics datasets

We collected major single-cell multi-omics sequencing datasets from a recent bench-marking study [64], comprising 5 single-cell RNA + protein datasets and 9 single-cell RNA + ATAC datasets. These datasets were generated using a variety of sequencing technologies, including CITE-seq[1], SHARE-seq[4], SNARE-seq[3], ISSAAC-seq[5], and 10x Multiome (https://www.10xgenomics.com/products/). To ensure high-quality data, we employed Seurat V4’s quality control filters with parameters outlined in the original publications, effectively filtering out low-quality cells, RNAs in scRNA-seq data, and peaks in scATAC-seq data. Further details can be found in Supplementary Tables 1, as well as the ‘Data availability’ section.

##### Spatial datasets

###### Slide-seqV2 mouse hippocampus

The SlideSeqV2 dataset [6], representing the mouse hippocampus, was retrieved using the datasets.slideseqv2() function from Squidpy(v1.6.1). This dataset comprises a hippocampal puck with 41,786 observations at near-cellular resolution and 4,000 genes. Since the dataset was preprocessed with log-transformed counts, we performed a reverse log normalization to derive the raw counts required for compatibility with Garfield. To facilitate benchmarking, we referenced the organizational annotations provided by the Allen Brain Atlas and manually annotated the dataset to generate ground truth for the experiment.

###### Mouse olfactory bulb (MOB) datasets

We collected two MOB datasets, sequenced using Stereo-seq and Slide-seqV2, respectively. The Slide-seqV2 dataset comprises 20,139 observations at near-cellular resolution and 21,220 genes, while the subcellular MOB dataset from Stereo-seq contains 19,109 observations and 27,106 genes. These two datasets were subsequently merged into a single AnnData object, containing 39,248 observations and 18,279 genes. Batch effects arising from different sequencing platforms will be corrected during model training to ensure data alignment.

###### High-resolution Xenium human breast cancer

The 10x Xenium human breast cancer dataset and its associated metadata were obtained from 10x Genomics (https://www.10xgenomics.com/products/xenium-in-situ/preview-dataset-human-breast). This dataset comprises a total of 282,363 cells across two replicates: 164,000 cells in replicate 1 and 118,363 cells in replicate 2, with 313 genes probed. To ensure data quality, we filtered out cells with fewer than 10 total counts across all genes or with non-zero counts in fewer than 3 distinct genes.

###### NanoString CosMx human non-small cell lung cancer(NSCLC)

The NanoString CosMx dataset for NSCLC [13] was sourced from https://nanostring.com/products/cosmx-spatial-molecular-imager/ffpe-dataset/nsclc-ffpe-dataset/. It encompasses 8 tissue sections derived from 5 donors, with a total of 800,327 cells distributed as follows: 93,206, 97,487, and 91,691 cells for replicate 1, replicate 2, and replicate 3 of donor 1(lung5); 83,723 cells from donor 2(lung6); 77,391 and 115,676 cells for replicates 1 and 2 of donor 3(lung9), respectively; 66,489 cells from donor 4(lung12); and 76,536 cells from donor 5(lung13). Of these datasets, all but donor 5 (lung13) were employed to construct a detailed spatial transcriptomic atlas, providing a comprehensive representation of NSCLC. Conversely, donor 5 (lung13) was reserved as query data to assess Garfield’s capacity for seamless and rapid integration of newly acquired datasets, highlighting its adaptability and robustness.

To ensure data quality, cells with fewer than 50 total counts were excluded, yielding 766,313 high-quality cells. During model training, we accounted for variability by incorporating covariates such as sample identity and donor origin.

###### Spatial ATAC-RNA multi-modal mouse brain

The Spatial epigenome–transcriptome sequencing dataset for the mouse brain was obtained from Zhang et al. [65]. This dataset features a brain tissue sample from a juvenile (P22) mouse, comprising 9,215 observations at spot-level resolution, with 22,914 genes and 121,068 peaks. To ensure high-quality data, we filtered out genes and peaks with counts detected in fewer than 46 cells (0.05% of all cells). Subsequently, we selected the 3,000 most highly variable genes and 10,000 most highly variable peaks, as identified by Scanpy. Finally, the two modality-specific AnnData objects were integrated into a single multimodal mdata object, which served as the input for Garfield’s analysis.

#### 5.3.2 Experiment setup and baseline methods

##### Benchmarking for single-cell multi-modal integration task

In the single-cell multi-modal integration task, we systematically compared the performance of Garfield with four algorithms optimized for RNA and ATAC integration and another four designed for RNA and ADT integration. This benchmark study encompasses six SOTA integrated algorithms. Similar to SeuratV4 and Multigrate, Garfield demonstrates versatility, enabling integration across different multimodal datasets (RNA + ATAC or RNA + ADT). The parameter configurations for these algorithms were determined based on the following guidelines:

###### Garfield

The final joint embeddings were obtained by making minimal parameter adjustments based on the specific single-cell multi-modal dataset. For RNA+ATAC datasets, the ‘sub_data_type’ parameter was set to RNA+ATAC, and the genome parameter was configured as either human or mouse depending on the dataset’s species origin. For RNA+ADT datasets, only the ‘sub_data_type’ parameter was modified to RNA+ADT. All other parameters were kept at their default values to ensure consistency across analyses. A specific example can be found through https://garfield-bio.readthedocs.io/en/latest/tutorial/03.10x_pbmc_paired_scMulti_analysis.html.

###### SeuratV4

To perform single-cell multi-modal vertical integration, we adhered to the tutorial available on the Seurat(v4.1.0) website via https://satijalab.org/seurat/articles/weighted_nearest_neighbor_analysis.

###### MultiVI

Vertical integration using MultiVI(scvi-tools v1.2.0) was conducted in accordance with the instructions provided in the https://docs.scvi-tools.org/en/stable/tutorials/notebooks/multimodal/MultiVI_tutorial.html.

###### MOFA+

We integrated single-cell RNA expression and chromatin accessibility data following the guidelines outlined in the MOFA’s website(v1.9.2) at https://muon-tutorials.readthedocs.io/en/latest/single-cellrna-atac/pbmc10k/3-Multimodal-Omics-Data-Integration.html.

###### totalVI

We conducted the analysis based on the instructions from the https://docs.scvi-tools.org/en/stable/tutorials/notebooks/multimodal/cite_scrna_integration_w_totalVI.html. Key parameters for the analysis were set as latent_distribution = normal and n_layers_decoder = 2.

###### scArches

We followed the guidelines provided on the scArches(v0.6.1) website for RNA+ADT integration: https://scarches.readthedocs.io/en/latest/totalvi_surgery_pipeline.html. The model was trained with parameters ‘epochs = 200, plan_kwargs = dict’.

###### Multigrate

Single-cell multi-omics data integration was performed using the original Multigrate model(v0.0.2), as described in the https://multigrate.readthedocs.io/en/latest/notebooks/paired_integration_cite-seq.html. Specifically, we employed the ‘nb’ loss function for RNA data and the ‘mse’ function for ATAC and ADT data.

##### Benchmarking for spatial niches discovery task

We compared Garfield(v1.0.0) to three other spatial transcriptome algorithms: NicheCompass (v0.2.1), CellCharter (v0.3.2), and GraphST (v1.1.1). For this benchmark study, we selected mouse hippocampus data, characterized by its well-defined tissue structure, as the data source. We conducted eight training runs for each method on this dataset, while varying the number of neighbors. The neighbor counts ranged from 4 to 16, incremented in steps of 4, with each configuration evaluated in two independent runs. Specifically, the parameter configurations for these competing algorithms were set as follows:

###### Garfield

Garfield models were trained using the Adam optimizer with an initial learning rate of 0.001. A learning rate scheduler was employed, reducing the learning rate by a factor of 0.1 after 4 epochs without improvement, while early stopping was applied with a patience of 8 epochs. The ‘GAT’ network was selected as the Garfield encoder, and the augment_type was set to ‘dropout’ for training and generating cell embeddings, with all other parameters left at their default settings. Detailed training configurations and procedures are available in the https://garfield-bio.readthedocs.io/en/latest/tutorial/05.Garfield_spatial_niche_slideseqv2_mouse_hippocampus.html.

###### NicheCompass

We followed the guidelines provided in the https://nichecompass.readthedocs.io/en/latest/tutorials/notebooks/mouse_cns_single_sample.html for model training, using the default hyperparameters.

###### CellCharter

As described in the https://cellcharter.readthedocs.io/en/latest/notebooks/codex_mouse_spleen.html, we utilized scVI (v1.2.1) for dimensionality reduction and data integration. CellCharter(v0.3.2) was then applied with default hyperparameters to extract latent representations.

###### GraphST

We adhered to https://deepst-tutorials.readthedocs.io/en/latest for model training, employing the default hyperparameters provided for GraphST(v1.1.1).

#### 5.3.3 Benchmark metrics

##### Single-cell integration task

For comprehensive benchmarking of single-cell multi-omics integration, we employed ARI[66], NMI[67], cASW[68], and cLISI[69] as key metrics. ARI and NMI were used as primary measures to evaluate the concordance between known cell-type labels and the clusters identified by the Leiden algorithm. To ensure fair comparisons across algorithms, we automatically determined an appropriate resolution for each algorithm based on the number of original cell types, ensuring a consistent number of clusters. The average silhouette width (ASW) metric assessed the precision of cell–cell distances calculated by each integration algorithm. Specifically, we used cell-type label-based ASW (cASW) to evaluate each algorithm’s ability to preserve biological variation. A higher cASW indicates better cell-type separation, reflecting improved accuracy in grouping cells within clusters (intra-cluster similarity) while maintaining distinctiveness from other clusters (inter-cluster dissimilarity).

We also utilized LISI to assess cell-type separation, denoting it as cLISI in our study. A lower cLISI value indicates more effective cell-type separation and greater conservation of biological variation. To ensure consistent interpretation of results, we applied linear transformations[70] so that higher cLISI values represent improved performance.

Finally, we introduced the biological variation conservation score, a total metric for evaluating integration algorithms. This score is calculated as the mean of ARI, NMI, cASW, and cLISI, providing a comprehensive measure of each algorithm’s ability to preserve biological variation across datasets.

##### Spatial integration task

For comprehensive benchmarking of spatial data integration, we employed ARI[66], NMI[67], AMI, HOM[71], ASW[68] as key metrics. These metrics were computed using the scikit-learn[72] or scIB[70] packages. To mitigate potential biases introduced by the resolution parameter in the Leiden algorithm, we performed Leiden clustering across a range of resolutions, from 0.1 to 2.0 in increments of 0.1, using the latent embedding from each algorithm. The performance of each algorithm was then evaluated based on the highest Adjusted Rand Index (ARI) and Normalized Mutual Information (NMI) values achieved across these resolutions.

###### Adjusted Rand Index (ARI)

The Adjusted Rand Index quantifies the agreement between clustering results and ground truth labels by comparing pairwise groupings. Let *C* denote the ground truth labels and *K* the predicted labels. Define:

- *m*: The number of sample pairs assigned to the same group in both *C* and *K*.
- *n*: The number of sample pairs assigned to different groups in both *C* and *K*.

The unadjusted Rand Index (RI) is calculated as:

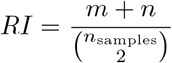

where 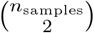 represents the total possible sample pairs.

Since RI is sensitive to random label assignments, particularly with similar numbers of clusters and samples, ARI adjusts for chance agreement by incorporating the expected value *E*[*RI*]. The adjusted formula is:

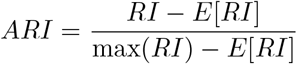

This adjustment ensures ARI scores close to zero for random clusterings, providing a robust and normalized measure for evaluating clustering performance.

###### Normalized Mutual Information (NMI)

NMI is s metric designed to quantify the alignment between predicted labels and the true labels. Given two sets of labels, *U* and *V*, representing the same *N* objects, the entropy of each set reflects the uncertainty associated with its partitions.

The entropy of *U* can be expressed as:

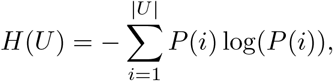

where *P* (*i*) denotes the probability of randomly selecting an object from *U* that belongs to partition *U*_*i*_, calculated as:

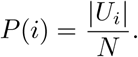

Similarly, the entropy of *V* is computed as:

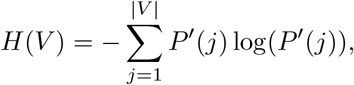

with *P* ^′^(*j*) representing the probability of an object being assigned to partition *V*_*j*_, defined as:

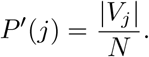

To measure the shared information between *U* and *V*, the mutual information (MI) is defined as:

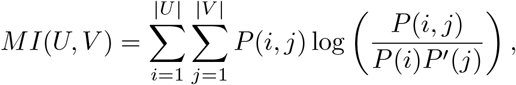

where *P* (*i, j*) is the joint probability that an object belongs to both partition *U*_*i*_ in *U* and *V*_*j*_ in *V*, given by:

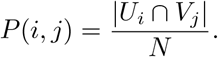

The normalized mutual information (NMI) provides a scale-invariant comparison by dividing MI by the average entropy of *U* and *V*, as follows:

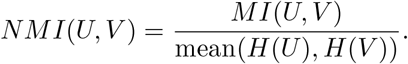

This normalization ensures that the score remains within a consistent range, facilitating comparisons across different datasets or clustering algorithms.

###### Adjusted Mutual Information (AMI)

The Adjusted Mutual Information (AMI) refines the Mutual Information (MI) by accounting for the expected similarity between two label assignments due to chance. It is defined as:

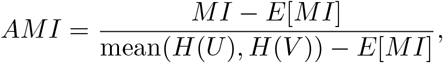

where *E*[*MI*] represents the expected value of the Mutual Information under random labelings.

The expected mutual information, *E*[*MI*(*U, V*)], is computed as:

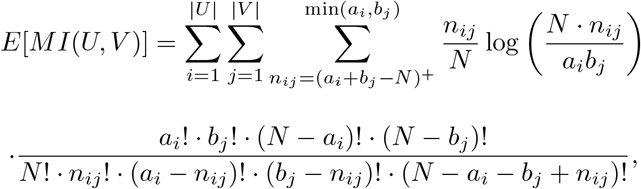

where *a*_*i*_ = |*U*_*i*_ |and *b*_*j*_ = |*V*_*j*_| denote the number of elements in partitions *U*_*i*_ and *V*_*j*_, respectively, and *N* is the total number of objects.

Here, *n*_*ij*_ represents the number of shared elements between partition *U*_*i*_ in *U* and *V*_*j*_ in *V*. The term (*a*_*i*_+*b*_*j*_ − *N*)^+^ ensures non-negative lower bounds for the summation.

By adjusting for *E*[*MI*], the AMI eliminates bias caused by chance, ensuring a robust and normalized comparison of clustering results.

###### Homogeneity (HOM)

Homogeneity is a clustering evaluation metric that assesses whether each cluster consists exclusively of members from a single class. The principle behind this metric is that a well-defined cluster should contain only data points that are similar to each other.

The homogeneity score ranges from 0 to 1, where:

- A score of 1 indicates perfect homogeneity, meaning every cluster contains members from only one class.
- A score of 0 signifies complete heterogeneity, where clusters are composed of members from multiple classes.

This metric provides a measure of how effectively a clustering algorithm preserves class integrity within clusters.

###### Average Silhouette Width (ASW)

The Average Silhouette Width (ASW) metric evaluates the precision of cell–cell distances computed by each integration algorithm. In our study, niche label-based ASW was employed to assess each algorithm’s effectiveness in preserving niche-specific biological variation.

### 5.4 Downstream analyses

#### Data Visualization

To visualize the latent representations generated by various methods, we employed the Uniform Manifold Approximation and Projection (UMAP) algorithm to reduce the latent matrices to two dimensions. A k-nearest neighbor graph was first constructed using `Scanpy’s pp.neighbors()` function with default configurations, followed by dimensionality reduction via `tl.umap()`. The resulting two-dimensional embeddings were then plotted through `scanpy.pl.umap()`. For spatial coordinate visualization, we used the `scanpy.pl.embedding()` function within the Scanpy(v1.9.1) package.

#### Niche Identification

Leiden clustering is conducted on the model’s latent space via the `scanpy.tl.leiden()` function, defining each cluster as a niche. Relationships among niches are analyzed through hierarchical clustering using `scanpy.tl.dendrogram()`, with niche labels as input and “ward” or “single” linkage methods applied.

#### Neighborhood Composition Analysis

The neighborhood composition score evaluates the spatial proximity between identified niches and cell types. We utilized the built-in calc_neighbor_prop function from the Garfield package to compute this score. Specifically, Garfield firstly computes the nearest neighbors of each cell using spatial coordinates, identifies the cell types within the neighborhood, and calculates the proportion of each cell type among the neighbors.

#### Spatial CHAOS Score Calculation

To evaluate the spatial continuity and compactness of niches identified by Garfield, we computed the spatial CHAOS score [46]. For each slice *s*, a one-nearest-neighbor (1NN) graph was constructed for all spatial locations within each niche *m*. The edge weight between locations *i* and *j* was defined as:

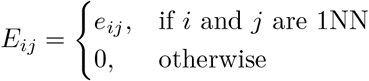

Here, *e*_*ij*_ represents the Euclidean distance between *i* and *j*. The spatial CHAOS score for slice *s* was calculated as:

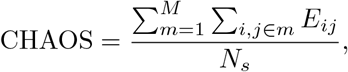

where *N*_*s*_ is the total number of spatial locations in slice *s, M* is the total number of spatial niches, and *i, j* ∈ *m* denotes that *i* and *j* belong to niche *m*.

#### Enrichment Score Calculation for Cell Types

To assess the enrichment of a specific cell type within a niche, we utilized an “enrichment ratio” metric. This ratio compares the observed number of cells of the given type to the expected number, assuming a uniform distribution of that cell type across all niches. The expected cell count within the niche was calculated using the Chi-square test. An enrichment ratio exceeding 1, combined with a P-value below 0.05, indicates significant enrichment of the cell type in the niche.

#### Proximal Cell-Cell Co-location Network Construction

To construct the proximal cell-cell interaction network, a spatial connectivity graph was created using the `squidpy.gr.spatial_neighbors()` function, with the interaction matrix computed via the `squidpy.gr.interaction_matrix()` function from the Squidpy(v1.6.1) package. The interaction strength *C*_*ij*_ between cell types *i* and *j* was calculated as: 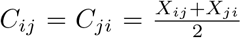,where *X*_*ij*_ and *X*_*ji*_ are the strengths of cell-cell interactions between cell types *i* and *j*.

To test statistical significance, a permutation-based test (default *n* = 1, 000) was performed by shuffling cell type labels while preserving the spatial connectivity graph. The *P* -value was determined as the proportion of permuted interaction strengths exceeding the true interaction strength. Cell types with fewer than 10 cells were excluded from the analysis.

#### Differential Expression Gene Analysis

After obtaining the niches labels, differential expressed gene (DEG) analysis was performed on the identified niches using built-in `calc_niches_marker` function from Garfield package to identify DEGs.

#### Functional Enrichment Analysis

After obtaining the DEGs via Differential expression gene analysis, functional enrichment analysis was performed using built-in `calc_niches_enrich` function from Garfield package to explore functional terms of each niche.

#### Cell–Cell Interaction Analysis

CellChat V2[49](v2.1.0) was employed to infer, analyze, and visualize intercellular communication between niches in spatial Xenium human breast cancer data. Ligand-receptor interactions were analyzed using ‘Scanpy’ normalized expression matrices, with interaction pairs sourced from CellChatDB, a curated database of mouse and human ligand-receptor interactions. Over-expressed ligands and receptors were identified for each niche, and communication probabilities were inferred by evaluating all ligand-receptor interactions linked to each signaling pathway.

## Supplementary information

**Fig. S1.**
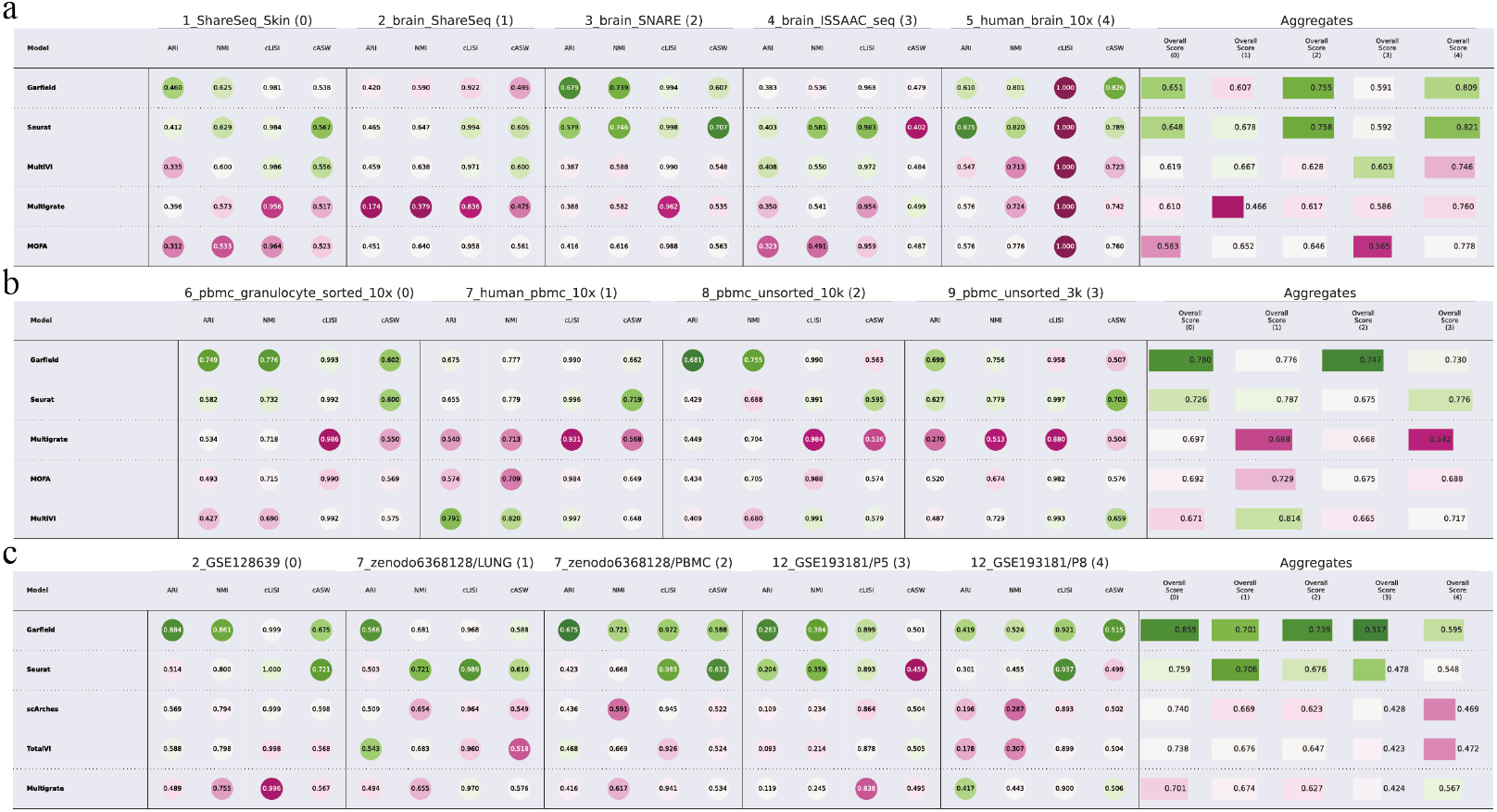
Comparative Benchmarking of Garfield Across Diverse Single-Cell Multi-Omics Datasets. **a**. Garfield’s performance is compared with Seurat V4, MultiVI, MOFA+, and Multigrate across five single-cell RNA+ATAC datasets. Metrics such as ARI, NMI, cLISI, and cASW demonstrate Garfield’s superior clustering accuracy, biological variation preservation. **b**. Similar to panel a, but using datasets derived from the remaining 4 single-cell RNA+ATAC datasets. **c**. Garfield is assessed on four single-cell RNA+ADT datasets (GSE128639, Zenodo_LUNG, Zenodo_PBMC, and GSE193181), comparing its performance to Seurat V4, scArches, TotalVI, and Multigrate.

**Fig. S2.**
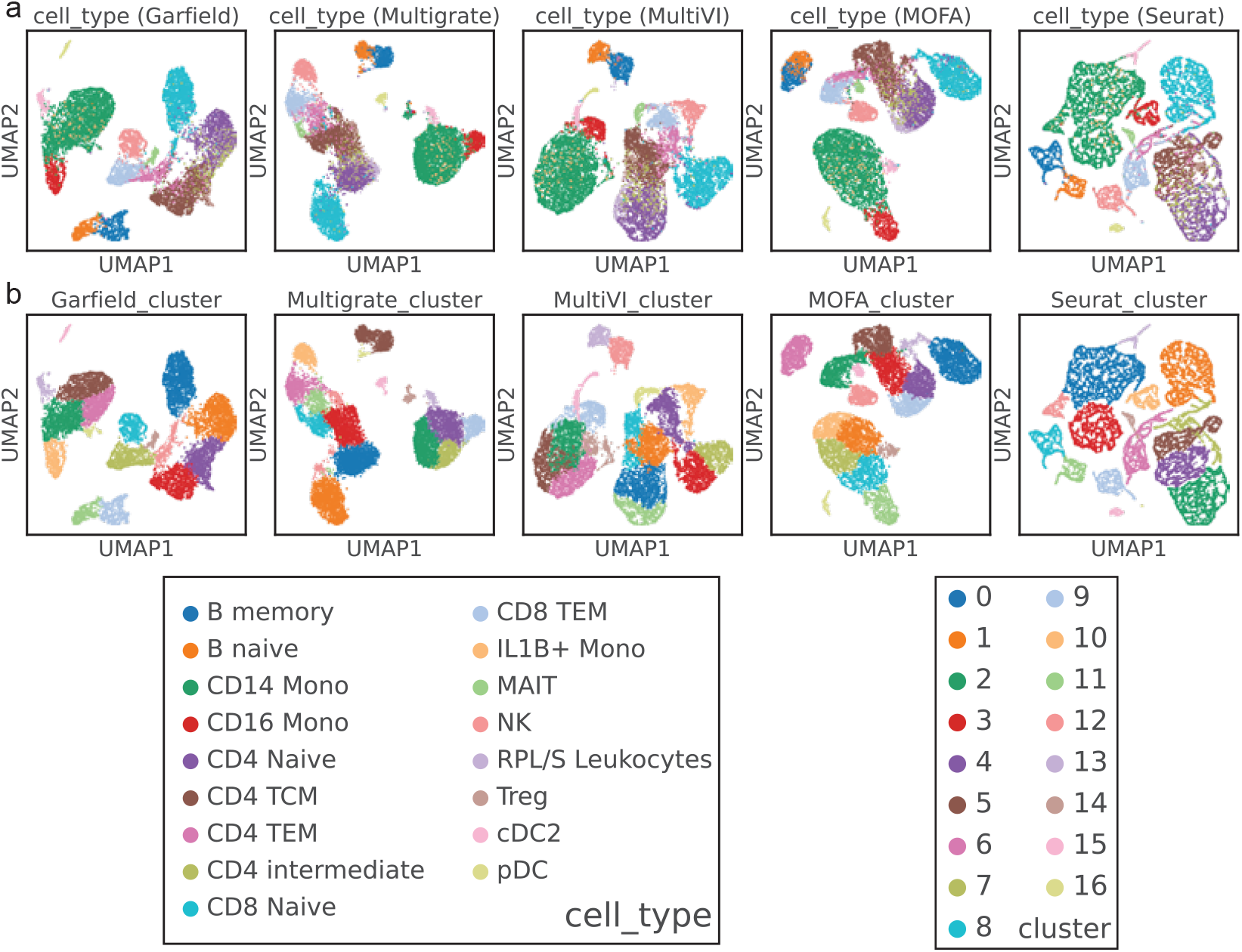
Garfield extracts cross-modality representations from parallel single-cell RNA-ATAC datasets. To simplify the presentation, we focus on the results from one representative dataset (*pbmc_granulocyte_sorted_10x*), as the results from other datasets exhibit similar patterns. **a**. The UMAP visualization displays latent representations obtained through different methods, with points colored by the original cell type labels. **b**. The UMAP visualization highlights latent representations derived from different methods, with points colored by molecular clusters identified using the different approaches.

**Fig. S3.**
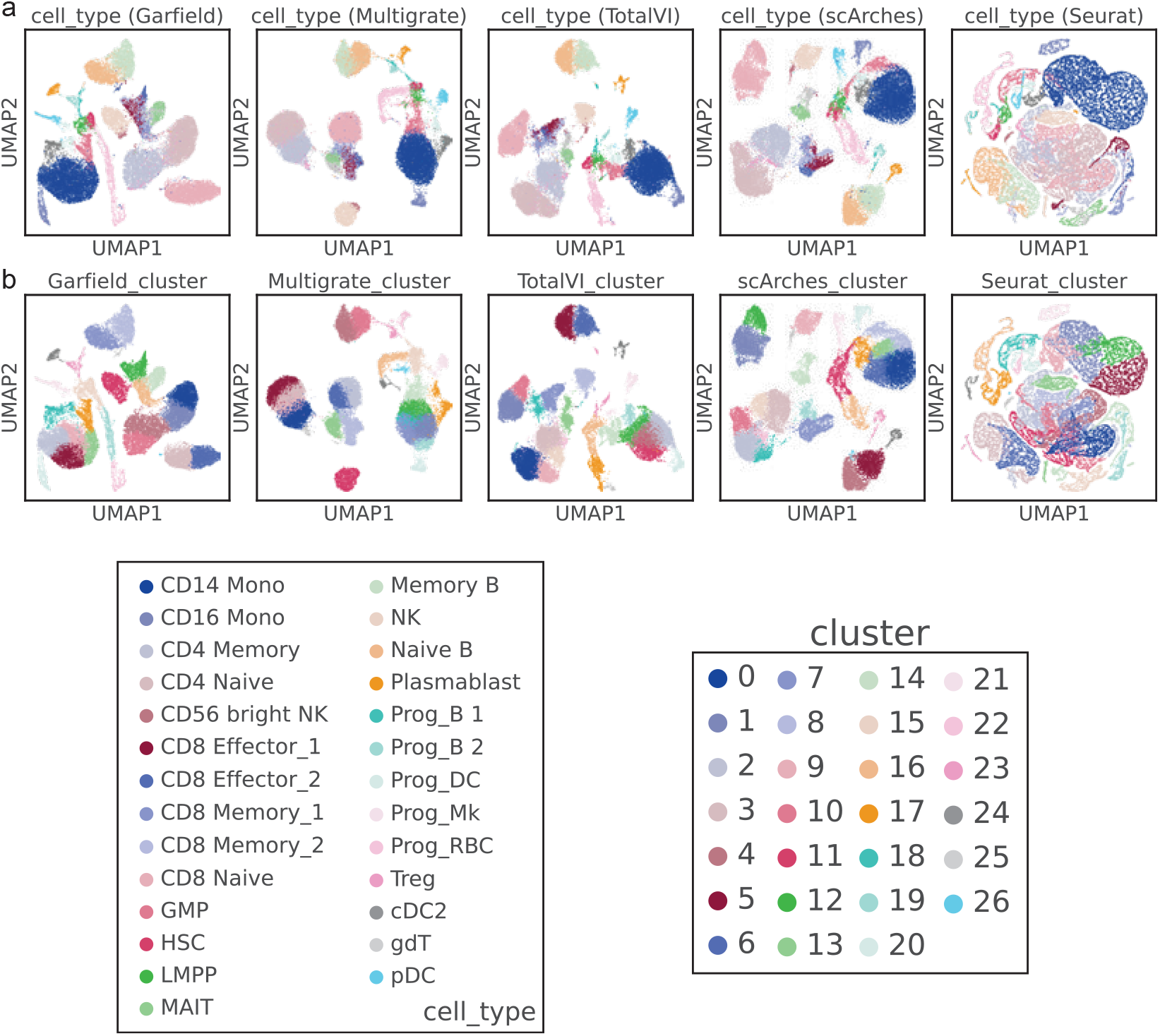
Garfield extracts cross-modality representations from parallel single-cell RNA– ADT datasets. For clarity, only the results from one representative dataset (GSE128639) are presented, as the outcomes from other datasets are analogous. **a**. The UMAP visualizations of the latent representations—colored by the original cell type labels—obtained using various methods. **b**. The UMAP visualizations of the latent representations—colored according to molecular clusters identified via different approaches.

**Fig. S4.**
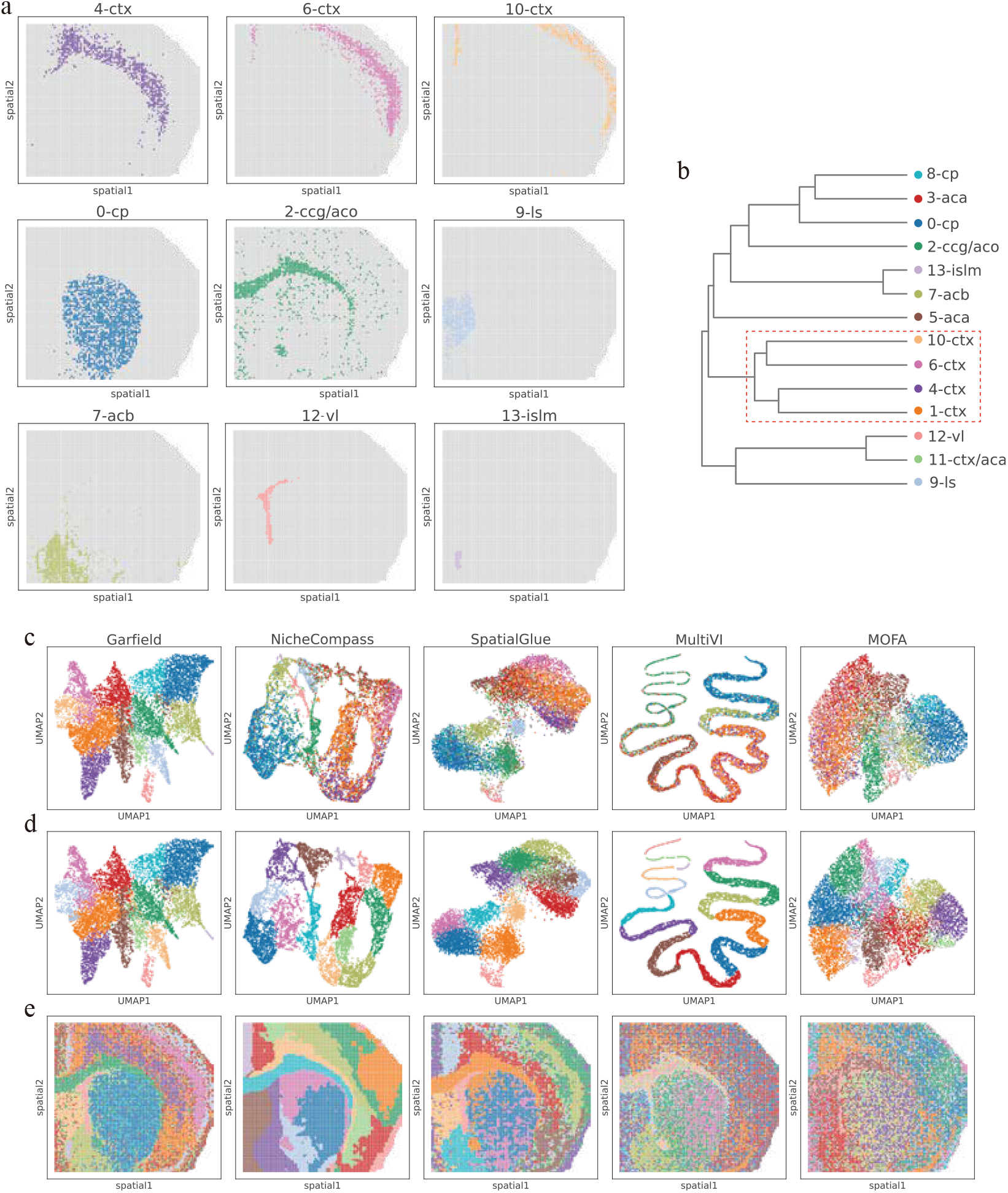
Garfield enables high-resolution dissection of spatial epigenome-transcriptome landscapes in mouse brain samples. **a**. Spatial expression patterns of primary niches (e.g., ctx, cp, ls), and each niche is marked by its spatial coordinates. **b**. Dendrogram illustrating the hierarchical relationships between niches based on Garfield latent. **c-e**. UMAP and spatial maps comparing niche detection results across Garfield and alternative methods (NicheCompass, SpatialGlue, MultiVI, MOFA), which colored by ground truth niches labels(c) and niches identified based on different methods (d-e).

**Fig. S5.**
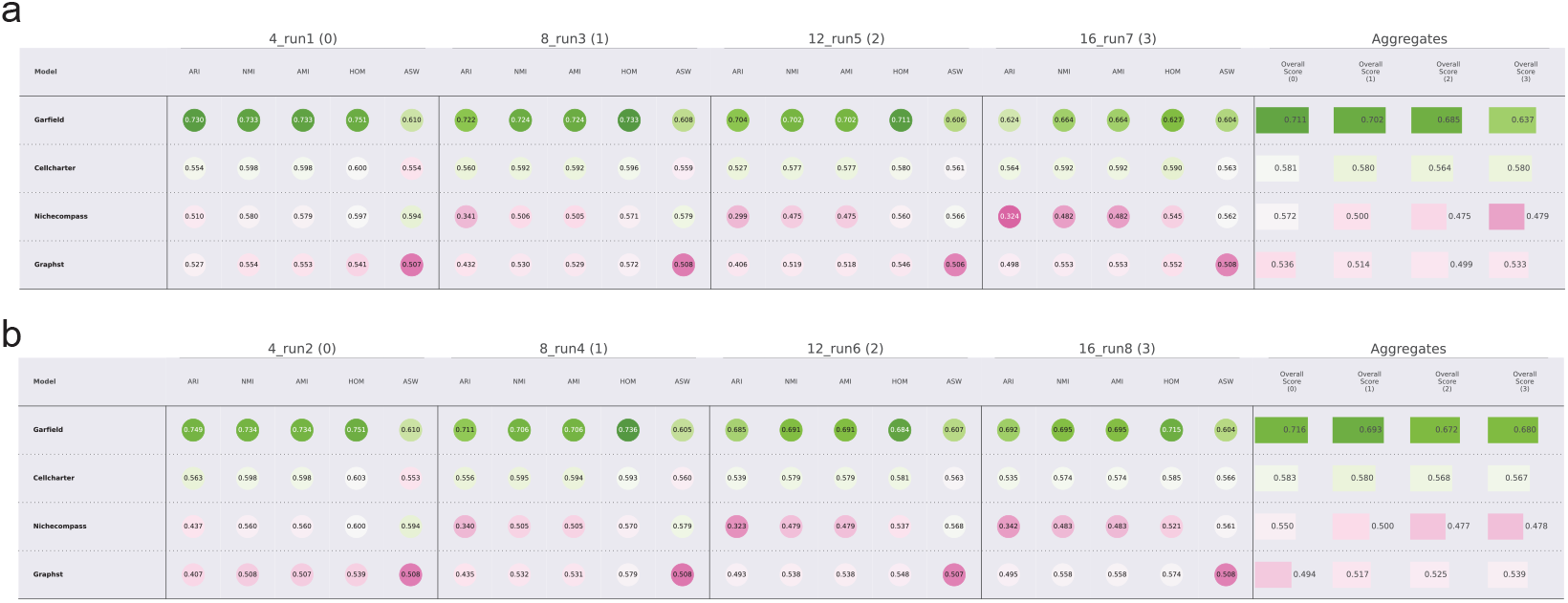
Benchmarking Garfield and related algorithms with varying numbers of neighbors or seeds on a spatial multi-omics dataset. **a**. Garfield consistently achieves the highest scores across evaluation metrics, including ARI, NMI, AMI, HOM, and ASW, demonstrating its reliability in preserving clustering accuracy, spatial organization, and biological variation across different neighbor sizes. **b**. The benchmarking assesses performance variation caused by random seed differences. Garfield remains the most stable and accurate across all metrics, significantly outperforming competing methods. Its aggregate scores reinforce its robustness in generating reproducible results under different initialization conditions.

**Fig. S6.**
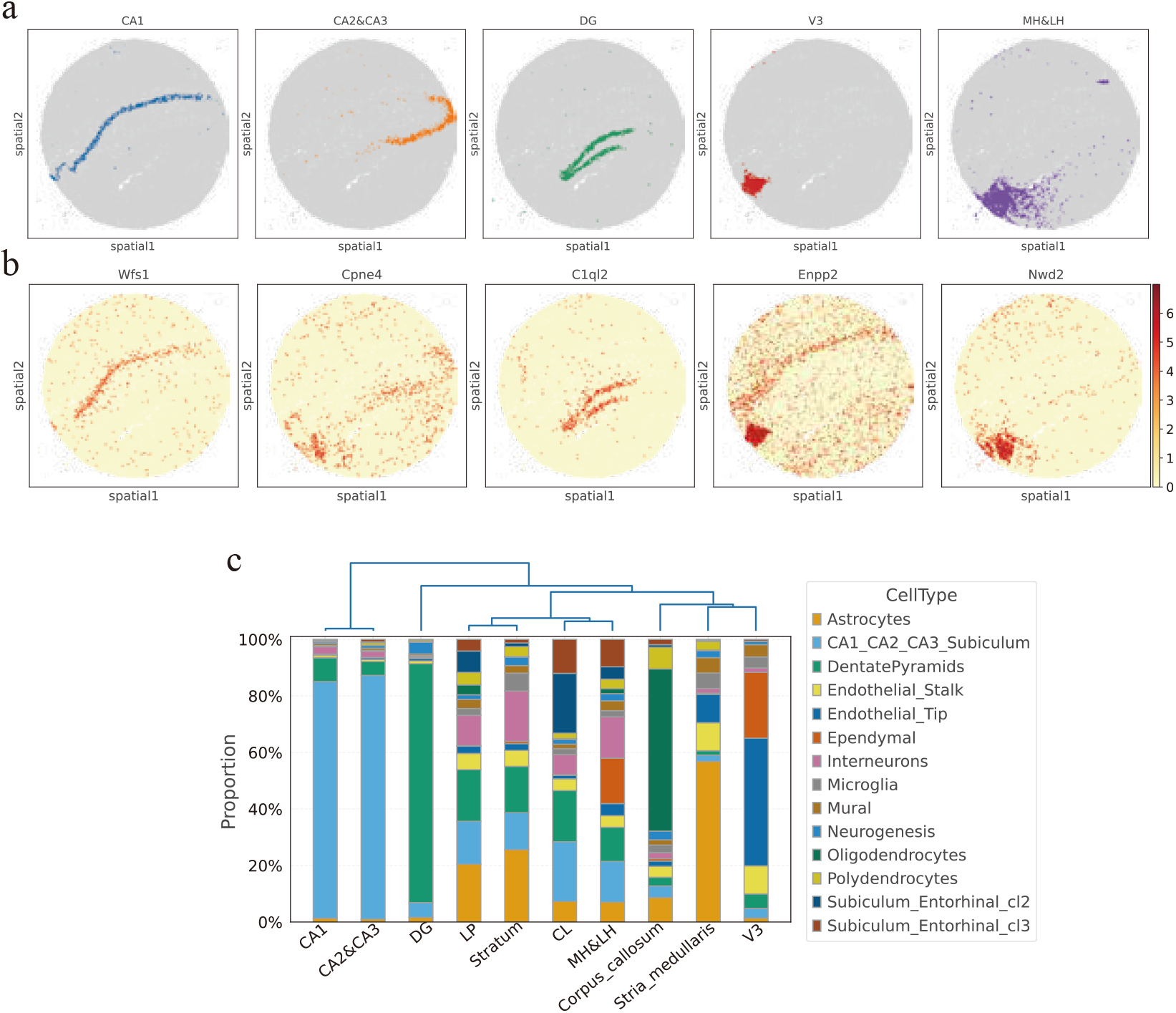
Analysis of heterogeneity of hippocampal tissue using Garfield. **a**. Visualization of spatial niches corresponding to major anatomical regions, including CA1, CA2&CA3, DG (dentate gyrus), V3, and MH&LH (medial and lateral habenula). **b**. Spatial expression of region-specific marker genes, including Wfs1 (CA1), Cpne4 (CA2&CA3), C1ql2 (DG), Enpp2 (V3), and Nwd2 (MH&LH). **c**. Proportional representation of major cell types across hippocampal regions, including astrocytes, interneurons, oligodendrocytes, endothelial cells (stalk and tip), microglia, and dentate pyramids. Hierarchical clustering shows similarities in cellular composition among niches.

**Fig. S7.**
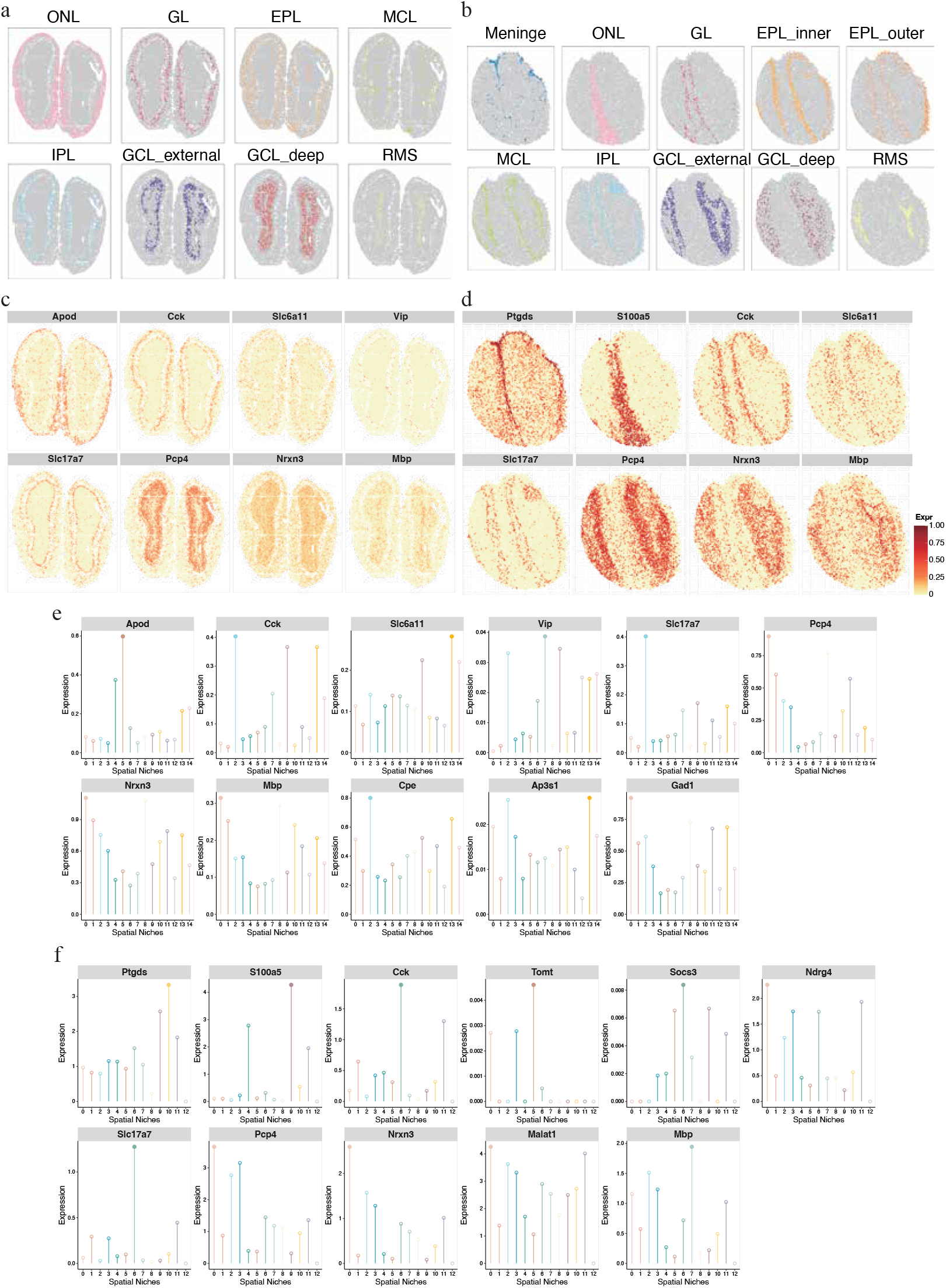
Analysis of heterogeneity of mouse olfactory bulb tissue from different sequencing technologies using Garfield. a–b. Scatter plots showing the spatial distribution of key MoB-related tissue structures in slices from Stereo-seq (a) and Slide-seqV2 (b). **c–d**. Scatter plots depicting the spatial pattern of key well-known markers in slices from Stereo-seq (c) and Slide-seqV2 (d). **e–f**. Lollipop plots illustrating the mean expression of key marker genes (y-axis) in slices from Stereo-seq (e) and Slide-seqV2 (f) across spatial inches identified by Garfield (x-axis), with solid circles highlighting the highest enrichment within each niche.

**Fig. S8.**
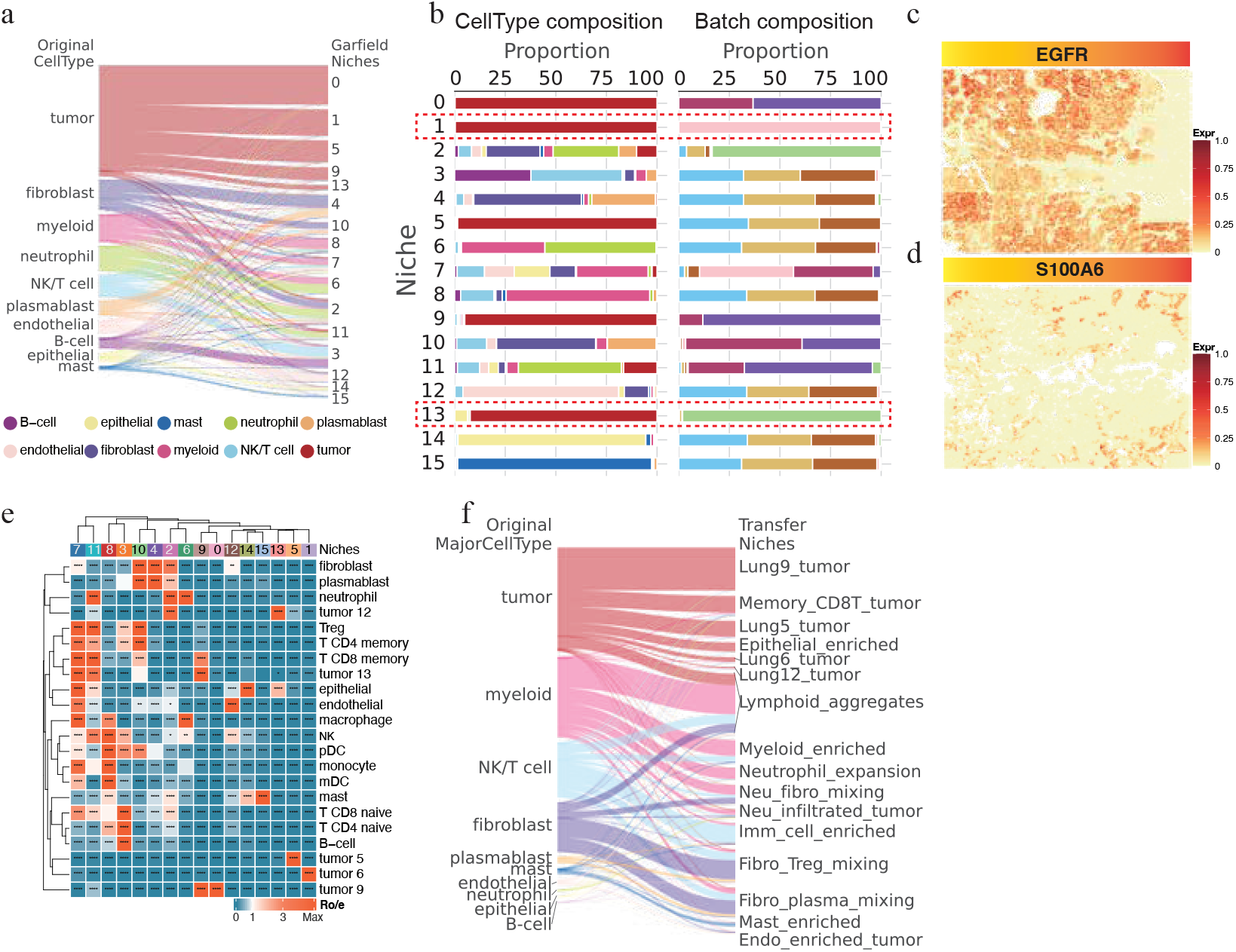
Analysis of Garfield niches reveals cellular heterogeneity and spatial organization in the NSCLC tumor microenvironment. **a**. Mapping of major cell types to Garfield-defined niches, visualizing the distribution and contribution of cell types within each niche. **b**. Proportions of major cell types(left) or data source(right) in different niches. **c-d**. Spatial expression maps of two hub genes (EGFR and S100A6) for Niche 1 and Niche 13, showing their distinct and niche-specific enrichment patterns. **e**. Preferential localization of each cell type in different niches, based on the ratio of observed to expected cell numbers (Ro/e). A ratio of 1 is indicated by white, with values >1 signifying enrichment in the specific niche. **f**. Sankey plot illustrates how the original tumor cell map to the transferred niches.

**Fig. S9.**
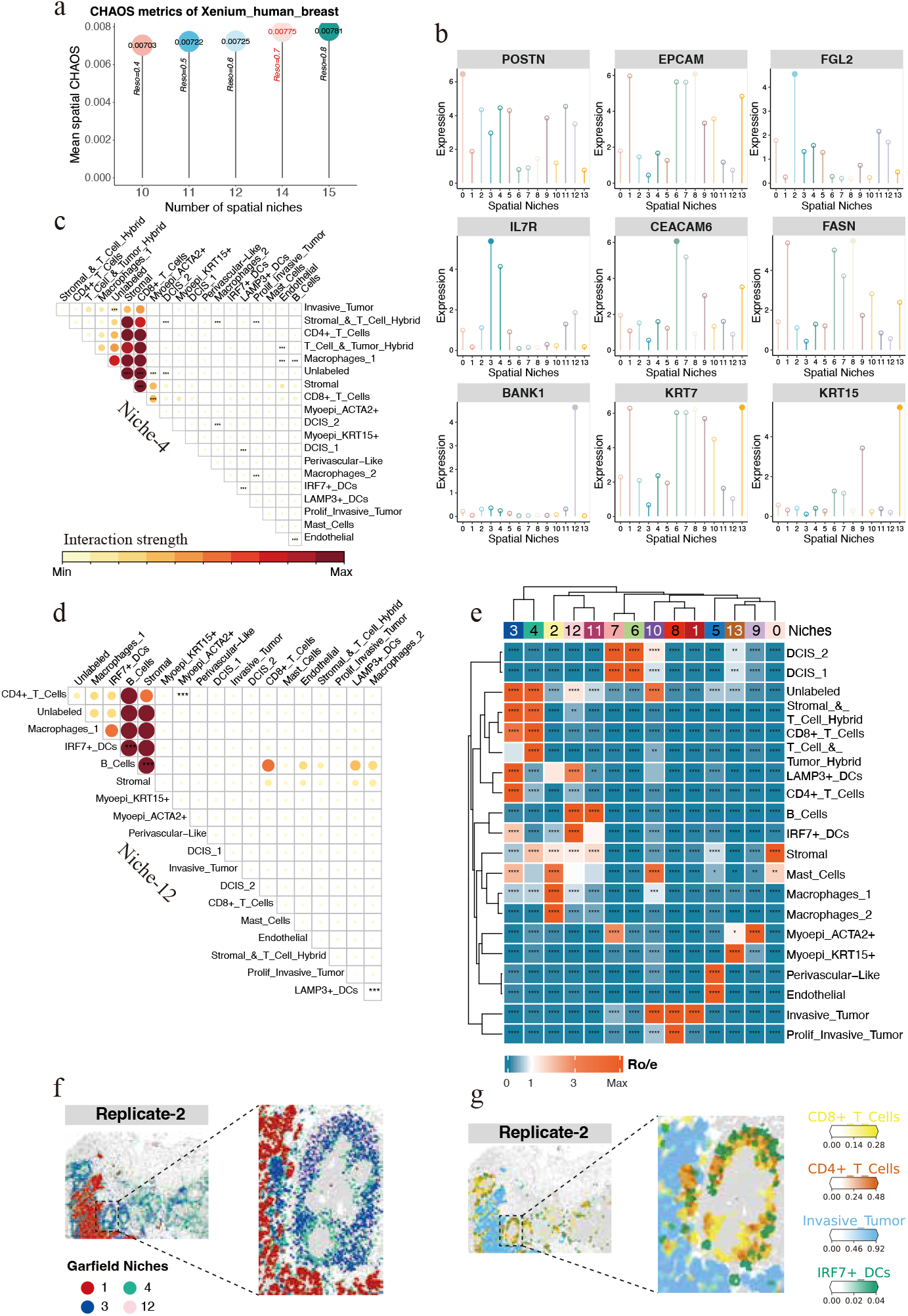
Garfield niche analysis uncovers cellular heterogeneity and spatial organization within the breast cancer microenvironment. **a**. Bar plots illustrating CHAOS values for BC tissue slices sequenced by 10X, showing the variation in pre-specified spatial niche resolution. The selected resolution (highlighted in red) was used for subsequent niche discovery. **b**. Lollipop plots showing the mean expression of key marker genes (y-axis) across spatial niches identified by Garfield (x-axis), with solid circles marking the highest enrichment in each niche. **c-d**. Heatmaps illustrating proximal cell-cell interactions within niche 4 (c), and niche 12 (d), with the color scale indicating the interaction strength between cell types i and j. **e**. Preferential localization of spatial niches identified by Garfield, based on the ratio of observed to expected cell numbers (Ro/e). A ratio of 1 is indicated by white, with values >1 signifying enrichment in the specific niche. **f**. Spatial distribution of tumor-specific niche and lymphoid niches within replicate-2. The proximal local network at the tumor edge is highlighted. **g**. Spatial distribution of invasive tumor cells (yellow), CD8+ T cells (red), CD4+ T cells (blue) and IRF7+ DCs (green) within the replicate-2 tissue sections. Gray regions represent background areas.

## Data availability

The datasets analyzed in this study are all from publicly available datasets (Supplementary Table 1). Specifically, for single-cell multi-omics datasets, they were obtained from a recent benchmarking study [64], comprising 5 single-cell RNA + protein datasets and 9 single-cell RNA + ATAC datasets. For spatial sequencing datasets, the SlideSeqV2 dataset, representing the mouse hippocampus, was retrieved using the datasets.slideseqv2() function from Squidpy. The mouse olfactory bulb tissue data generated by Stereoseq and Slide-seqV2 platforms can be accessed from https://github.com/JinmiaoChenLab/SEDR_analyses and https://singlecell.broadinstitute.org/single_cell/study/SCP815, respectively. The 10x Xenium human breast cancer dataset and its associated metadata were obtained from 10x Genomics (https://www.10xgenomics.com/products/xenium-in-situ/preview-dataset-human-breast). The NanoString CosMx dataset for NSCLC was sourced from https://nanostring.com/products/cosmx-spatial-molecular-imager/ffpe-dataset/nsclc-ffpe-dataset/. The Spatial epigenome–transcriptome sequencing mouse brain dataset was retrieved from https://www.ncbi.nlm.nih.gov/geo/query/acc.cgi?acc=GSE205055 (gene expression counts and spatial coordinates) and https://brain-spatial-omics.cells.ucsc.edu/ (peak counts and cell type labels). The details of all datasets used are available in the Methods. The annotation images from the Allen Mouse Brain Atlas can be accessed at https://mouse.brain-map.org/static/atlas.

## Code availability

Garfield is available as a Python package, maintained at https://github.com/zhou-1314/Garfield. The Jupyter notebooks for reproducing our analyses and benchmarking results are available at https://github.com/zhou-1314/Garfield-reproducibility. Documentation is provided at https://garfield-bio.readthedocs.io/.

## Acknowledgements

We thank X.F. and other authors for assistance in improving Garfield, X.L. for comments on the manuscript. We acknowledge the support of the Data Science Platform of Guangzhou National Laboratory and the Bio-medical Big Data Operating System (Bio-OS). This work has been supported by the Major Project of Guangzhou National Laboratory Grant No. GZNL2024A03001.

## Author contribution

W.Z., L.T., W.J. conceived and supervised the project. W.Z. designed and developed the Garfield algorithm. W.Z., and L.T. wrote the manuscript. W.Z., X.F., L.L., J.Z., performed the data analysis. W.Z. prepared the figures. W.J. and X.L. provided valuable suggestions. All authors read and approved the final paper.

## Appendix A Section title of first appendix

An appendix contains supplementary information that is not an essential part of the text itself but which may be helpful in providing a more comprehensive understanding of the research problem or it is information that is too cumbersome to be included in the body of the paper.

## References

[1] Stoeckius, M., Hafemeister, C., Stephenson, W., Houck-Loomis, B., Chattopadhyay, P.K., Swerdlow, H., Satija, R., Smibert, P.: Simultaneous epitope and transcriptome measurement in single cells. Nature methods 14(9), 865–868 (2017)

[2] Peterson, V.M., Zhang, K.X., Kumar, N., Wong, J., Li, L., Wilson, D.C., Moore, R., McClanahan, T.K., Sadekova, S., Klappenbach, J.A.: Multiplexed quantification of proteins and transcripts in single cells. Nature biotechnology 35(10), 936–939 (2017)

[3] Chen, S., Lake, B.B., Zhang, K.: High-throughput sequencing of the transcriptome and chromatin accessibility in the same cell. Nature biotechnology 37(12), 1452–1457 (2019)

[4] Ma, S., Zhang, B., LaFave, L.M., Earl, A.S., Chiang, Z., Hu, Y., Ding, J., Brack, A., Kartha, V.K., Tay, T., et al.: Chromatin potential identified by shared single-cell profiling of rna and chromatin. Cell 183(4), 1103–1116 (2020)

[5] Xu, W., Yang, W., Zhang, Y., Chen, Y., Hong, N., Zhang, Q., Wang, X., Hu, Y., Song, K., Jin, W., et al.: Issaac-seq enables sensitive and flexible multimodal profiling of chromatin accessibility and gene expression in single cells. Nature Methods 19(10), 1243–1249 (2022)

[6] Stickels, R.R., Murray, E., Kumar, P., Li, J., Marshall, J.L., Di Bella, D.J., Arlotta, P., Macosko, E.Z., Chen, F.: Highly sensitive spatial transcriptomics at near-cellular resolution with slide-seqv2. Nature biotechnology 39(3), 313–319 (2021)

[7] Chen, A., Liao, S., Cheng, M., Ma, K., Wu, L., Lai, Y., Qiu, X., Yang, J., Xu, J., Hao, S., et al.: Spatiotemporal transcriptomic atlas of mouse organogenesis using dna nanoball-patterned arrays. Cell 185(10), 1777–1792 (2022)

[8] Liu, Y., Yang, M., Deng, Y., Su, G., Enninful, A., Guo, C.C., Tebaldi, T., Zhang, D., Kim, D., Bai, Z., et al.: High-spatial-resolution multi-omics sequencing via deterministic barcoding in tissue. Cell 183(6), 1665–1681 (2020)

[9] Fu, X., Sun, L., Dong, R., Chen, J.Y., Silakit, R., Condon, L.F., Lin, Y., Lin, S., Palmiter, R.D., Gu, L.: Polony gels enable amplifiable dna stamping and spatial transcriptomics of chronic pain. Cell 185(24), 4621–4633 (2022)

[10] Cho, C.-S., Xi, J., Si, Y., Park, S.-R., Hsu, J.-E., Kim, M., Jun, G., Kang, H.M., Lee, J.H.: Microscopic examination of spatial transcriptome using seq-scope. Cell 184(13), 3559–3572 (2021)

[11] Lubeck, E., Coskun, A.F., Zhiyentayev, T., Ahmad, M., Cai, L.: Single-cell in situ rna profiling by sequential hybridization. Nature methods 11(4), 360–361 (2014)

[12] Chen, K.H., Boettiger, A.N., Móitt, J.R., Wang, S., Zhuang, X.: Spatially resolved, highly multiplexed rna profiling in single cells. Science 348(6233), 6090 (2015)

[13] He, S., Bhatt, R., Brown, C., Brown, E.A., Buhr, D.L., Chantranuvatana, K., Danaher, P., Dunaway, D., Garrison, R.G., Geiss, G., et al.: High-plex imaging of rna and proteins at subcellular resolution in fixed tissue by spatial molecular imaging. Nature Biotechnology 40(12), 1794–1806 (2022)

[14] Argelaguet, R., Cuomo, A.S., Stegle, O., Marioni, J.C.: Computational principles and challenges in single-cell data integration. Nature biotechnology 39(10), 1202– 1215 (2021)

[15] Stuart, T., Butler, A., Hoffman, P., Hafemeister, C., Papalexi, E., Mauck, W.M., Hao, Y., Stoeckius, M., Smibert, P., Satija, R.: Comprehensive integration of single-cell data. cell 177(7), 1888–1902 (2019)

[16] Ashuach, T., Gabitto, M.I., Koodli, R.V., Saldi, G.-A., Jordan, M.I., Yosef, N.: Multivi: deep generative model for the integration of multimodal data. Nature Methods 20(8), 1222–1231 (2023)

[17] Lotfollahi, M., Litinetskaya, A., Theis, F.J.: Multigrate: single-cell multi-omic data integration. BioRxiv, 2022–03 (2022)

[18] Argelaguet, R., Arnol, D., Bredikhin, D., Deloro, Y., Velten, B., Marioni, J.C., Stegle, O.: Mofa+: a statistical framework for comprehensive integration of multi-modal single-cell data. Genome biology 21, 1–17 (2020)

[19] Hao, Y., Hao, S., Andersen-Nissen, E., Mauck, W.M., Zheng, S., Butler, A., Lee, M.J., Wilk, A.J., Darby, C., Zager, M., et al.: Integrated analysis of multimodal single-cell data. Cell 184(13), 3573–3587 (2021)

[20] Long, Y., Ang, K.S., Li, M., Chong, K.L.K., Sethi, R., Zhong, C., Xu, H., Ong, Z., Sachaphibulkij, K., Chen, A., et al.: Spatially informed clustering, integration, and deconvolution of spatial transcriptomics with graphst. Nature Communications 14(1), 1155 (2023)

[21] Dong, K., Zhang, S.: Deciphering spatial domains from spatially resolved transcriptomics with an adaptive graph attention auto-encoder. Nature communications 13(1), 1739 (2022)

[22] Varrone, M., Tavernari, D., Santamaria-Martínez, A., Walsh, L.A., Ciriello, G.: Cellcharter reveals spatial cell niches associated with tissue remodeling and cell plasticity. Nature Genetics 56(1), 74–84 (2024)

[23] Li, Y., Zhang, J., Gao, X., Zhang, Q.C.: Tissue module discovery in single-cell-resolution spatial transcriptomics data via cell-cell interaction-aware cell embedding. Cell Systems (2024)

[24] Birk, S., et al.: Quantitative characterization of cell niches in spatial atlases. bioRxiv (2024) 10.1101/2024.02.21.581428

[25] Kipf, T.N., Welling, M.: Variational graph auto-encoders. arXiv preprint 1611.07308 (2016)

[26] Lotfollahi, M., Naghipourfar, M., Luecken, M.D., Khajavi, M., Büttner, M., Wagenstetter, M., Avsec, Ž., Gayoso, A., Yosef, N., Interlandi, M., et al.: Mapping single-cell data to reference atlases by transfer learning. Nature biotechnology 40(1), 121–130 (2022)

[27] Wolf, F.A., Angerer, P., Theis, F.J.: Scanpy: large-scale single-cell gene expression data analysis. Genome biology 19, 1–5 (2018)

[28] Palla, G., Spitzer, H., Klein, M., Fischer, D., Schaar, A.C., Kuemmerle, L.B., Rybakov, S., Ibarra, I.L., Holmberg, O., Virshup, I., et al.: Squidpy: a scalable framework for spatial omics analysis. Nature methods 19(2), 171–178 (2022)

[29] Wang, J., Ma, A., Chang, Y., Gong, J., Jiang, Y., Qi, R., Wang, C., Fu, H., Ma, Q., Xu, D.: scgnn is a novel graph neural network framework for single-cell rna-seq analyses. Nature communications 12(1), 1882 (2021)

[30] Brody, S., Alon, U., Yahav, E.: How attentive are graph attention networks? arXiv preprint 2105.14491 (2021)

[31] Veličković, P., Cucurull, G., Casanova, A., Romero, A., Lio, P., Bengio, Y.: Graph attention networks. arXiv preprint 1710.10903 (2017)

[32] Kipf, T.N., Welling, M.: Semi-supervised classification with graph convolutional networks. arXiv preprint 1609.02907 (2016)

[33] Cai, X., Huang, C., Xia, L., Ren, X.: Lightgcl: Simple yet effective graph contrastive learning for recommendation. arXiv preprint 2302.08191 (2023)

[34] Gretton, A., Borgwardt, K.M., Rasch, M.J., Schölkopf, B., Smola, A.: A kernel two-sample test. The Journal of Machine Learning Research 13(1), 723–773 (2012)

[35] Lotfollahi, M., Naghipourfar, M., Theis, F.J., Wolf, F.A.: Conditional out-of-sample generation for unpaired data using trvae. arXiv preprint 1910.01791 (2019)

[36] Kartha, V.K., Duarte, F.M., Hu, Y., Ma, S., Chew, J.G., Lareau, C.A., Earl, A., Burkett, Z.D., Kohlway, A.S., Lebofsky, R., et al.: Functional inference of gene regulation using single-cell multi-omics. Cell genomics 2(9) (2022)

[37] Li, C., Virgilio, M.C., Collins, K.L., Welch, J.D.: Multi-omic single-cell velocity models epigenome–transcriptome interactions and improves cell fate prediction. Nature biotechnology 41(3), 387–398 (2023)

[38] La Manno, G., Soldatov, R., Zeisel, A., Braun, E., Hochgerner, H., Petukhov, V., Lidschreiber, K., Kastriti, M.E., Lönnerberg, P., Furlan, A., et al.: Rna velocity of single cells. Nature 560(7719), 494–498 (2018)

[39] Moses, L., Pachter, L.: Museum of spatial transcriptomics. Nature methods 19(5), 534–546 (2022)

[40] You, Y., Fu, Y., Li, L., Zhang, Z., Jia, S., Lu, S., Ren, W., Liu, Y., Xu, Y., Liu, X., et al.: Systematic comparison of sequencing-based spatial transcriptomic methods. Nature Methods 21(9), 1743–1754 (2024)

[41] Wang, Q., Ding, S.-L., Li, Y., Royall, J., Feng, D., Lesnar, P., Graddis, N., Naeemi, M., Facer, B., Ho, A., et al.: The allen mouse brain common coordinate framework: a 3d reference atlas. Cell 181(4), 936–953 (2020)

[42] Xu, H., Fu, H., Long, Y., Ang, K.S., Sethi, R., Chong, K., Li, M., Uddamvathanak, R., Lee, H.K., Ling, J., et al.: Unsupervised spatially embedded deep representation of spatial transcriptomics. Genome Medicine 16(1), 12 (2024)

[43] DeSisto, J., O’Rourke, R., Jones, H.E., Pawlikowski, B., Malek, A.D., Bonney, S., Guimiot, F., Jones, K.L., Siegenthaler, J.A.: Single-cell transcriptomic analyses of the developing meninges reveal meningeal fibroblast diversity and function. Developmental cell 54(1), 43–59 (2020)

[44] Gudjohnsen, S.A., Atacho, D.A., Gesbert, F., Raposo, G., Hurbain, I., Larue, L., Steingrimsson, E., Petersen, P.H.: Meningeal melanocytes in the mouse: distribution and dependence on mitf. Frontiers in Neuroanatomy 9, 149 (2015)

[45] Tepe, B., Hill, M.C., Pekarek, B.T., Hunt, P.J., Martin, T.J., Martin, J.F., Arenkiel, B.R.: Single-cell rna-seq of mouse olfactory bulb reveals cellular heterogeneity and activity-dependent molecular census of adult-born neurons. Cell reports 25(10), 2689–2703 (2018)

[46] Ma, Y., Zhou, X.: Accurate and efficient integrative reference-informed spatial domain detection for spatial transcriptomics. Nature Methods, 1–14 (2024)

[47] Zhou, X., Dong, K., Zhang, S.: Integrating spatial transcriptomics data across different conditions, technologies and developmental stages. Nature Computational Science 3(10), 894–906 (2023)

[48] Pandey, P.R., Saidou, J., Watabe, K.: Role of myoepithelial cells in breast tumor progression. Frontiers in bioscience: a journal and virtual library 15, 226 (2010)

[49] Jin, S., Plikus, M.V., Nie, Q.: Cellchat for systematic analysis of cell–cell communication from single-cell transcriptomics. Nature Protocols 20(1), 180–219 (2025)

[50] Long, Y., Ang, K.S., Sethi, R., Liao, S., Heng, Y., Olst, L., Ye, S., Zhong, C., Xu, H., Zhang, D., et al.: Deciphering spatial domains from spatial multi-omics with spatialglue. Nature Methods, 1–10 (2024)

[51] Chen, S., Zhu, B., Huang, S., Hickey, J.W., Lin, K.Z., Snyder, M., Greenleaf, W.J., Nolan, G.P., Zhang, N.R., Ma, Z.: Integration of spatial and single-cell data across modalities with weakly linked features. Nature Biotechnology 42(7), 1096–1106 (2024)

[52] Zhu, B., Chen, S., Bai, Y., Chen, H., Liao, G., Mukherjee, N., Vazquez, G., McIlwain, D.R., Tzankov, A., Lee, I.T., et al.: Robust single-cell matching and multimodal analysis using shared and distinct features. Nature Methods 20(2), 304–315 (2023)

[53] Chen, H., Ryu, J., Vinyard, M.E., Lerer, A., Pinello, L.: Simba: single-cell embedding along with features. Nature Methods 21(6), 1003–1013 (2024)

[54] Rong, Y., Huang, W., Xu, T., Huang, J.: Dropedge: Towards deep graph convolutional networks on node classification. arXiv preprint 1907.10903 (2019)

[55] Halko, N., Martinsson, P.-G., Tropp, J.A.: Finding structure with randomness: Probabilistic algorithms for constructing approximate matrix decompositions. SIAM review 53(2), 217–288 (2011)

[56] Hamilton, W., Ying, Z., Leskovec, J.: Inductive representation learning on large graphs. Advances in neural information processing systems 30 (2017)

[57] Zhang, X., Wang, X., Shivashankar, G., Uhler, C.: Graph-based autoencoder integrates spatial transcriptomics with chromatin images and identifies joint biomarkers for alzheimer’s disease. Nature Communications 13(1), 7480 (2022)

[58] Li, W., Yang, F., Wang, F., Rong, Y., Liu, L., Wu, B., Zhang, H., Yao, J.: scprotein: a versatile deep graph contrastive learning framework for single-cell proteomics embedding. Nature Methods 21(4), 623–634 (2024)

[59] Tschannen, M., Djolonga, J., Rubenstein, P.K., Gelly, S., Lucic, M.: On mutual information maximization for representation learning. arXiv preprint 1907.13625 (2019)

[60] Chen, T., Kornblith, S., Norouzi, M., Hinton, G.: A simple framework for contrastive learning of visual representations. In: International Conference on Machine Learning, pp. 1597–1607 (2020). PMLR

[61] Oord, A.v.d., Li, Y., Vinyals, O.: Representation learning with contrastive predictive coding. arXiv preprint 1807.03748 (2018)

[62] Li, Y., Hu, P., Liu, Z., Peng, D., Zhou, J.T., Peng, X.: Contrastive clustering. In: Proceedings of the AAAI Conference on Artificial Intelligence, vol. 35, pp. 8547–8555 (2021)

[63] Hu, W., Miyato, T., Tokui, S., Matsumoto, E., Sugiyama, M.: Learning discrete representations via information maximizing self-augmented training. In: International Conference on Machine Learning, pp. 1558–1567 (2017). PMLR

[64] Hu, Y., Wan, S., Luo, Y., Li, Y., Wu, T., Deng, W., Jiang, C., Jiang, S., Zhang, Y., Liu, N., et al.: Benchmarking algorithms for single-cell multi-omics prediction and integration. Nature Methods, 1–13 (2024)

[65] Zhang, D., Deng, Y., Kukanja, P., Agirre, E., Bartosovic, M., Dong, M., Ma, C., Ma, S., Su, G., Bao, S., et al.: Spatial epigenome–transcriptome co-profiling of mammalian tissues. Nature 616(7955), 113–122 (2023)

[66] Hubert, L., Arabie, P.: Comparing partitions. Journal of classification 2, 193–218 (1985)

[67] Strehl, A., Ghosh, J.: Cluster ensembles—a knowledge reuse framework for combining multiple partitions. Journal of machine learning research 3(Dec), 583–617 (2002)

[68] Rousseeuw, P.J.: Silhouettes: a graphical aid to the interpretation and validation of cluster analysis. Journal of computational and applied mathematics 20, 53–65 (1987)

[69] Korsunsky, I., Millard, N., Fan, J., Slowikowski, K., Zhang, F., Wei, K., Baglaenko, Y., Brenner, M., Loh, P.-r., Raychaudhuri, S.: Fast, sensitive and accurate integration of single-cell data with harmony. Nature methods 16(12), 1289–1296 (2019)

[70] Luecken, M.D., Büttner, M., Chaichoompu, K., Danese, A., Interlandi, M., Müller, M.F., Strobl, D.C., Zappia, L., Dugas, M., Colomé-Tatché, M., et al.: Benchmarking atlas-level data integration in single-cell genomics. Nature methods 19(1), 41–50 (2022)

[71] Rosenberg, A., Hirschberg, J.: V-measure: A conditional entropy-based external cluster evaluation measure. In: Proceedings of the 2007 Joint Conference on Empirical Methods in Natural Language Processing and Computational Natural Language Learning (EMNLP-CoNLL), pp. 410–420 (2007)

[72] Pedregosa, F., Varoquaux, G., Gramfort, A., Michel, V., Thirion, B., Grisel, O., Blondel, M., Prettenhofer, P., Weiss, R., Dubourg, V., et al.: Scikit-learn: Machine learning in python. the Journal of machine Learning research 12, 2825–2830 (2011)

