## Supplementary material for "Graph-based Contrastive Learning Enables Unified Integration and Niche Transfer Across Single-Cell and Spatial Multi-Omics": Fig.S1

a

| Model | 1_ShareSeq_Skin (0) | | | | 2_brain_ShareSeq (1) | | | | 3_brain_SNARE (2) | | | | 4_brain_ISSAAC_seq (3) | | | | 5_human_brain_10x (4) | | | | Aggregates | | | | |
|---|---|---|---|---|---|---|---|---|---|---|---|---|---|---|---|---|---|---|---|---|---|---|---|---|---|
| | ARI | NMI | cLISI | cASW | ARI | NMI | cLISI | cASW | ARI | NMI | cLISI | cASW | ARI | NMI | cLISI | cASW | ARI | NMI | cLISI | cASW | Overall Score (0) | Overall Score (1) | Overall Score (2) | Overall Score (3) | Overall Score (4) |
| Garfield | 0.460 | 0.625 | 0.981 | 0.538 | 0.420 | 0.590 | 0.922 | 0.495 | 0.679 | 0.739 | 0.994 | 0.607 | 0.383 | 0.536 | 0.968 | 0.479 | 0.610 | 0.801 | 1.000 | 0.826 | 0.651 | 0.607 | 0.755 | 0.591 | 0.809 |
| Seurat | 0.412 | 0.629 | 0.984 | 0.567 | 0.465 | 0.647 | 0.994 | 0.605 | 0.579 | 0.746 | 0.998 | 0.707 | 0.403 | 0.581 | 0.983 | 0.402 | 0.675 | 0.820 | 1.000 | 0.789 | 0.648 | 0.678 | 0.758 | 0.592 | 0.821 |
| MultiVI | 0.335 | 0.600 | 0.986 | 0.556 | 0.459 | 0.638 | 0.971 | 0.600 | 0.387 | 0.588 | 0.990 | 0.548 | 0.408 | 0.550 | 0.972 | 0.484 | 0.547 | 0.713 | 1.000 | 0.723 | 0.619 | 0.667 | 0.628 | 0.603 | 0.746 |
| Multigrate | 0.396 | 0.573 | 0.956 | 0.517 | 0.174 | 0.379 | 0.836 | 0.475 | 0.388 | 0.582 | 0.962 | 0.535 | 0.350 | 0.541 | 0.954 | 0.499 | 0.576 | 0.724 | 1.000 | 0.742 | 0.610 | 0.466 | 0.617 | 0.586 | 0.760 |
| MOFA | 0.312 | 0.533 | 0.964 | 0.523 | 0.451 | 0.640 | 0.958 | 0.561 | 0.416 | 0.616 | 0.988 | 0.563 | 0.323 | 0.491 | 0.959 | 0.487 | 0.576 | 0.776 | 1.000 | 0.760 | 0.583 | 0.652 | 0.646 | 0.565 | 0.778 |

b

| Model | 6_pbmc_granulocyte_sorted_10x (0) | | | | 7_human_pbmc_10x (1) | | | | 8_pbmc_unsorted_10k (2) | | | | 9_pbmc_unsorted_3k (3) | | | | Aggregates | | | |
|---|---|---|---|---|---|---|---|---|---|---|---|---|---|---|---|---|---|---|---|---|
| | ARI | NMI | cLISI | cASW | ARI | NMI | cLISI | cASW | ARI | NMI | cLISI | cASW | ARI | NMI | cLISI | cASW | Overall Score (0) | Overall Score (1) | Overall Score (2) | Overall Score (3) |
| Garfield | 0.749 | 0.776 | 0.993 | 0.602 | 0.675 | 0.777 | 0.990 | 0.662 | 0.681 | 0.755 | 0.990 | 0.563 | 0.699 | 0.756 | 0.958 | 0.507 | 0.780 | 0.776 | 0.747 | 0.730 |
| Seurat | 0.582 | 0.732 | 0.992 | 0.600 | 0.655 | 0.779 | 0.996 | 0.719 | 0.429 | 0.688 | 0.991 | 0.595 | 0.627 | 0.779 | 0.997 | 0.703 | 0.726 | 0.787 | 0.675 | 0.776 |
| Multigrate | 0.534 | 0.718 | 0.986 | 0.550 | 0.540 | 0.713 | 0.931 | 0.568 | 0.449 | 0.704 | 0.984 | 0.536 | 0.270 | 0.513 | 0.880 | 0.504 | 0.697 | 0.688 | 0.668 | 0.542 |
| MOFA | 0.493 | 0.715 | 0.990 | 0.569 | 0.574 | 0.709 | 0.984 | 0.649 | 0.434 | 0.705 | 0.988 | 0.574 | 0.520 | 0.674 | 0.982 | 0.576 | 0.692 | 0.729 | 0.675 | 0.688 |
| MultiVI | 0.427 | 0.690 | 0.992 | 0.575 | 0.791 | 0.820 | 0.997 | 0.648 | 0.409 | 0.680 | 0.991 | 0.579 | 0.487 | 0.729 | 0.993 | 0.659 | 0.671 | 0.814 | 0.665 | 0.717 |

c

| Model | 2_GSE128639 (0) | | | | 7_zenodo6368128/LUNG (1) | | | | 7_zenodo6368128/PBMC (2) | | | | 12_GSE193181/P5 (3) | | | | 12_GSE193181/P8 (4) | | | | Aggregates | | | | |
|---|---|---|---|---|---|---|---|---|---|---|---|---|---|---|---|---|---|---|---|---|---|---|---|---|---|
| | ARI | NMI | cLISI | cASW | ARI | NMI | cLISI | cASW | ARI | NMI | cLISI | cASW | ARI | NMI | cLISI | cASW | ARI | NMI | cLISI | cASW | Overall Score (0) | Overall Score (1) | Overall Score (2) | Overall Score (3) | Overall Score (4) |
| Garfield | 0.884 | 0.861 | 0.999 | 0.675 | 0.566 | 0.681 | 0.968 | 0.588 | 0.675 | 0.721 | 0.972 | 0.588 | 0.283 | 0.384 | 0.899 | 0.501 | 0.419 | 0.524 | 0.921 | 0.515 | 0.855 | 0.701 | 0.739 | 0.517 | 0.595 |
| Seurat | 0.514 | 0.800 | 1.000 | 0.721 | 0.503 | 0.721 | 0.989 | 0.610 | 0.423 | 0.668 | 0.985 | 0.631 | 0.204 | 0.359 | 0.893 | 0.458 | 0.301 | 0.455 | 0.937 | 0.499 | 0.759 | 0.706 | 0.676 | 0.478 | 0.548 |
| scArches | 0.569 | 0.794 | 0.999 | 0.598 | 0.509 | 0.654 | 0.964 | 0.549 | 0.436 | 0.591 | 0.945 | 0.522 | 0.109 | 0.234 | 0.864 | 0.504 | 0.196 | 0.287 | 0.893 | 0.502 | 0.740 | 0.669 | 0.623 | 0.428 | 0.469 |
| TotalVI | 0.588 | 0.798 | 0.998 | 0.568 | 0.543 | 0.683 | 0.960 | 0.518 | 0.468 | 0.669 | 0.926 | 0.524 | 0.093 | 0.214 | 0.878 | 0.505 | 0.178 | 0.307 | 0.899 | 0.504 | 0.738 | 0.676 | 0.647 | 0.423 | 0.472 |
| Multigrate | 0.489 | 0.755 | 0.996 | 0.567 | 0.494 | 0.655 | 0.970 | 0.576 | 0.416 | 0.617 | 0.941 | 0.534 | 0.119 | 0.245 | 0.838 | 0.495 | 0.417 | 0.443 | 0.900 | 0.506 | 0.701 | 0.674 | 0.627 | 0.424 | 0.567 |
