## Supplementary figures and images for "Graph-based Contrastive Learning Enables Unified Integration and Niche Transfer Across Single-Cell and Spatial Multi-Omics"

### Fig.S2

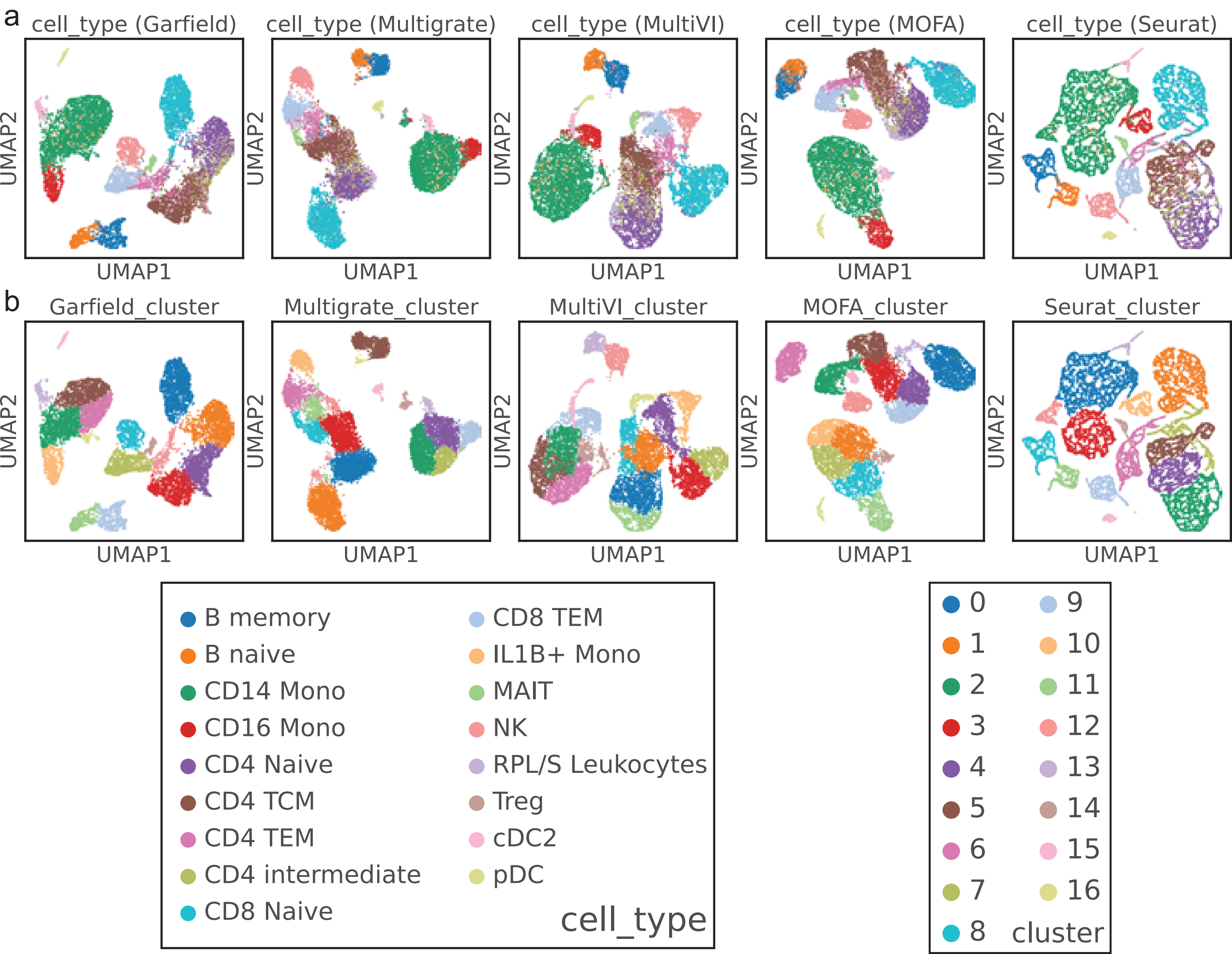

### Fig.S3

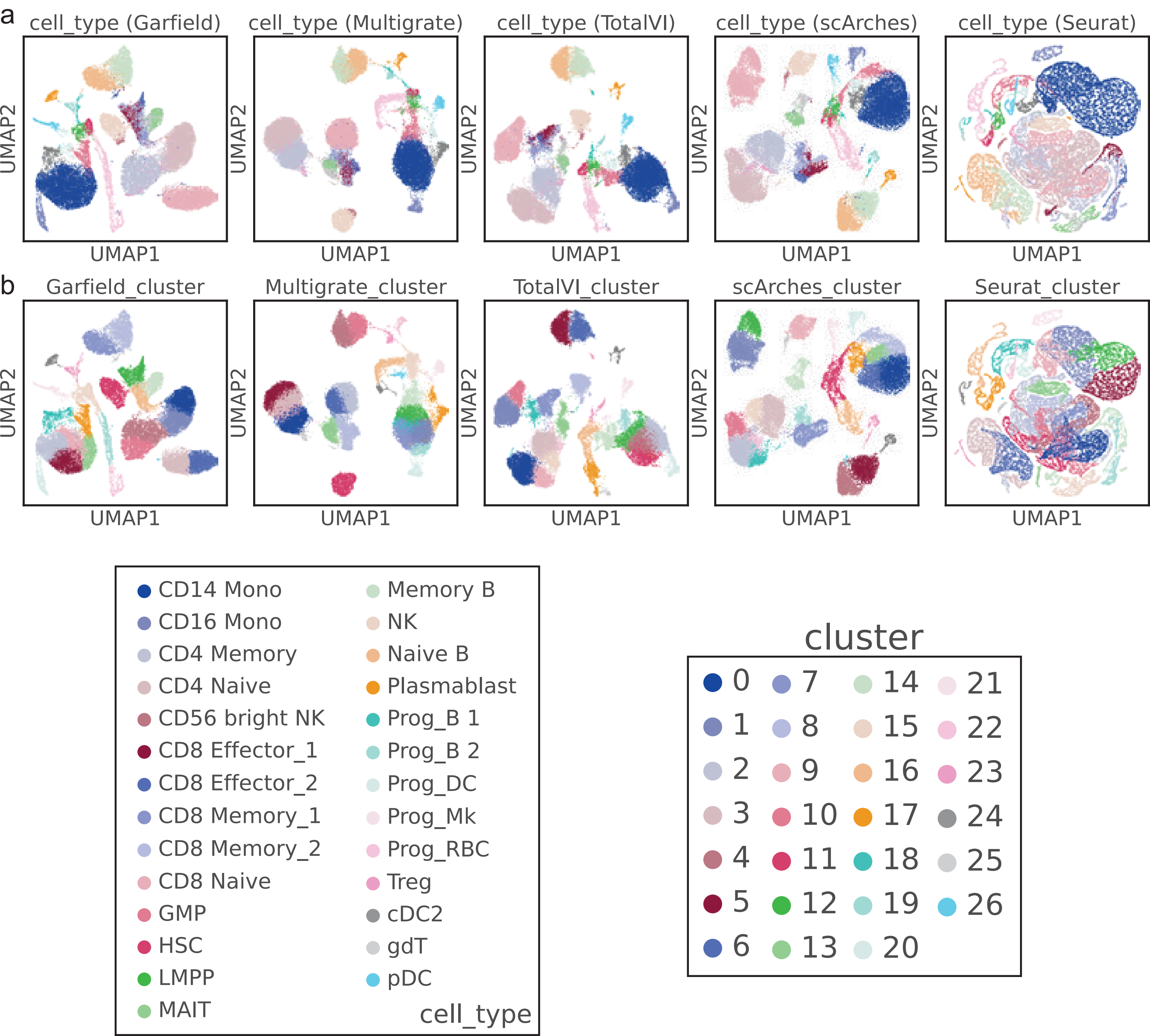

### Fig.S4

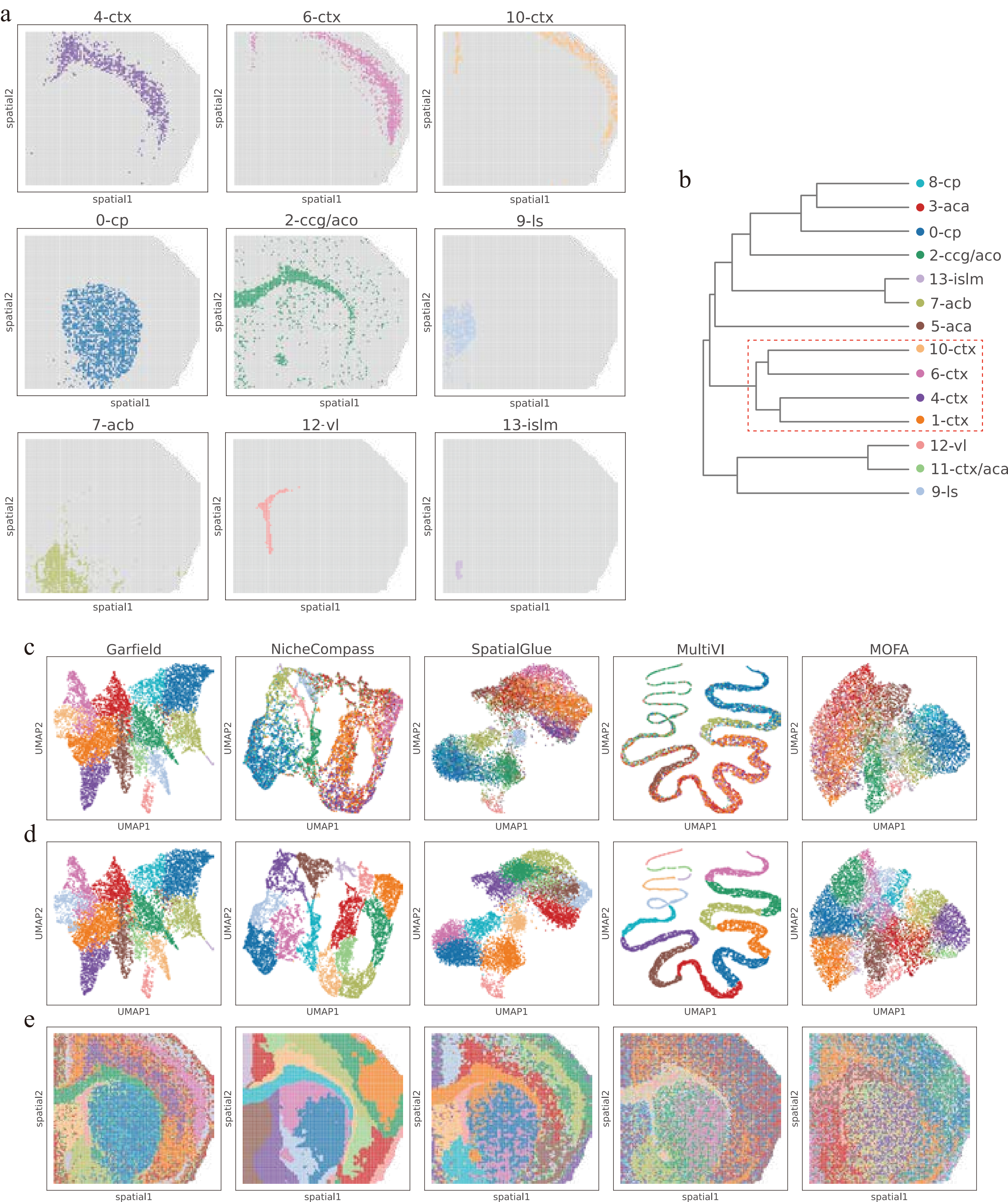

### Fig.S6

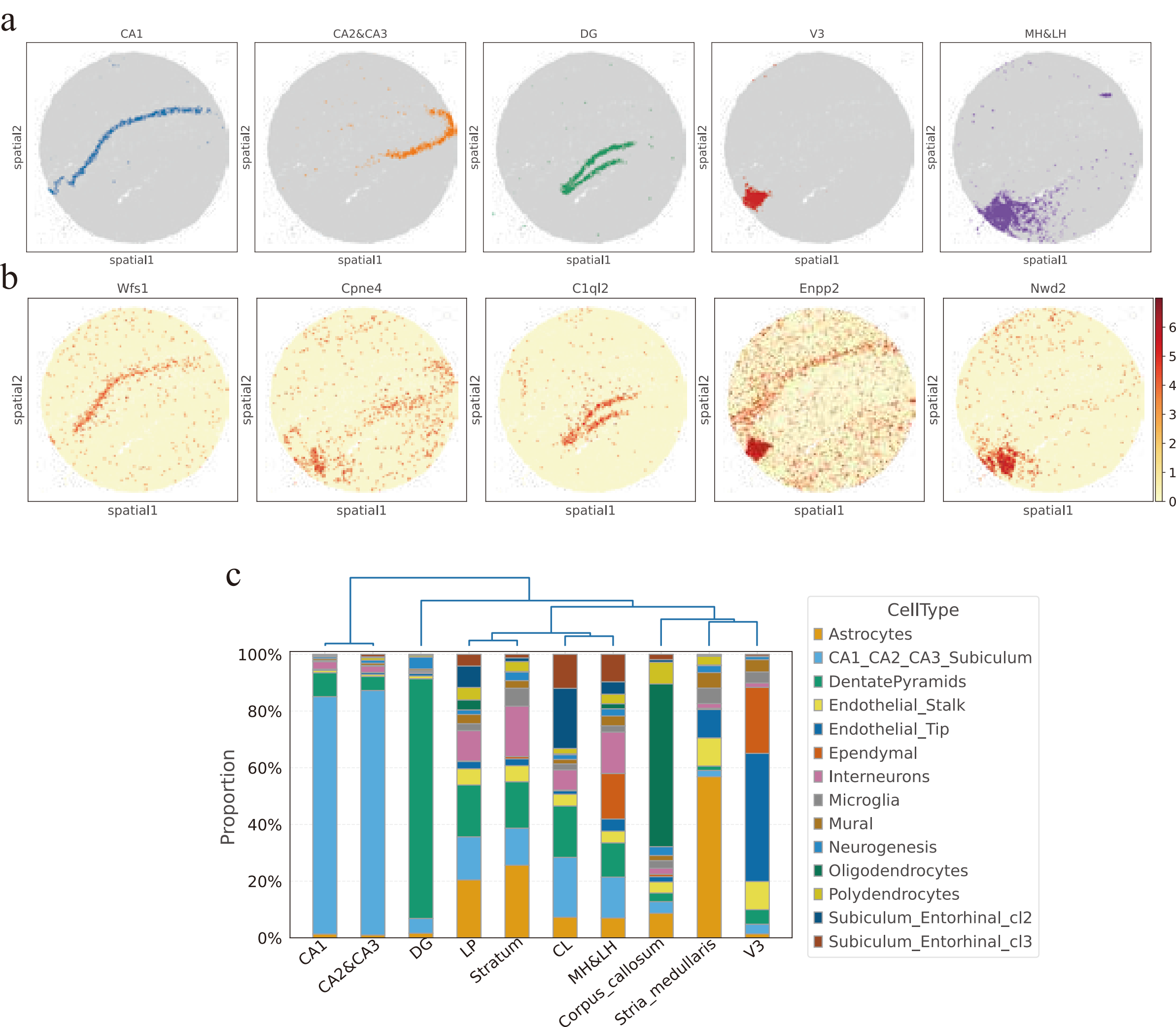

### Fig.S7

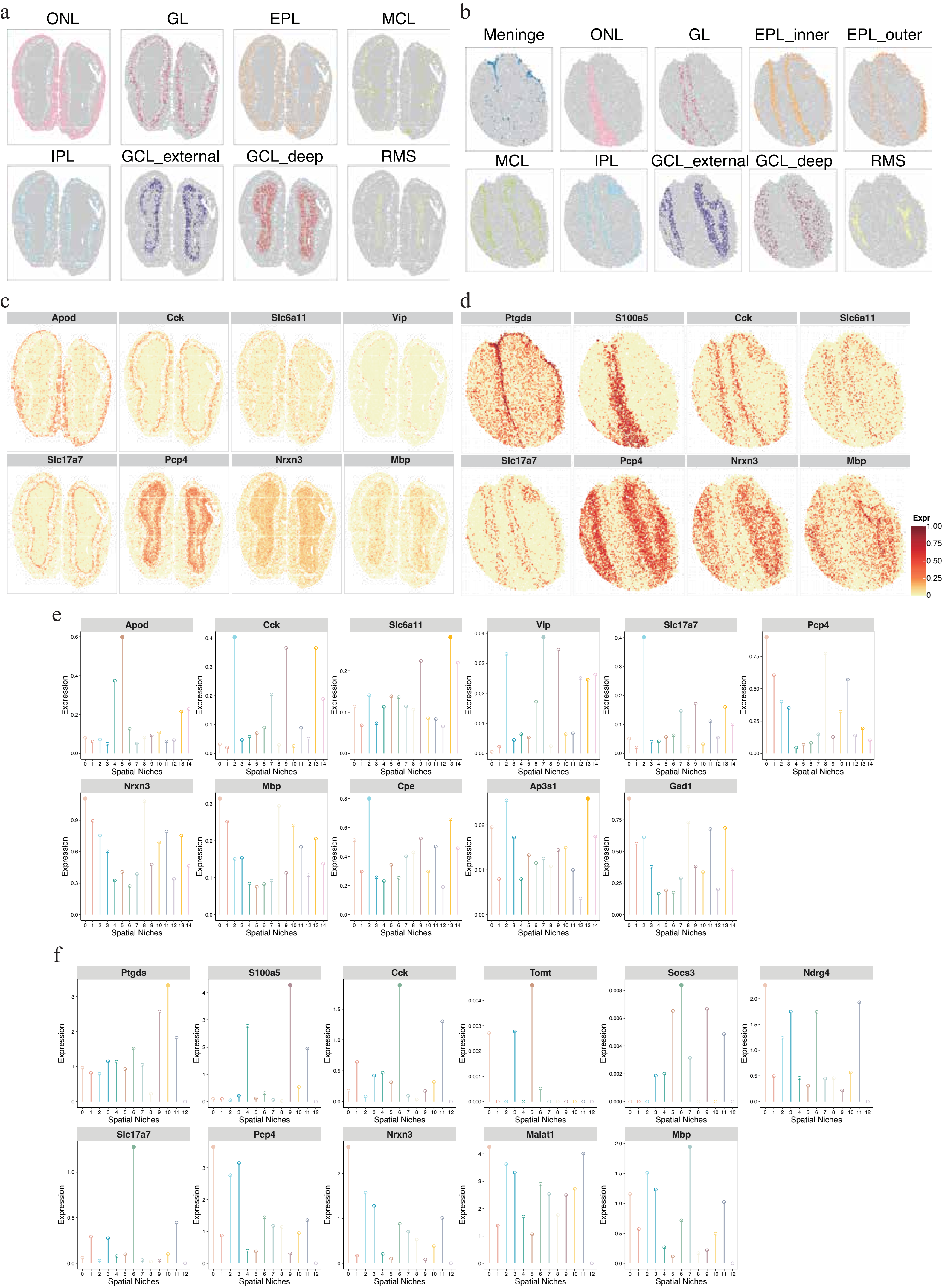

### Fig.S8

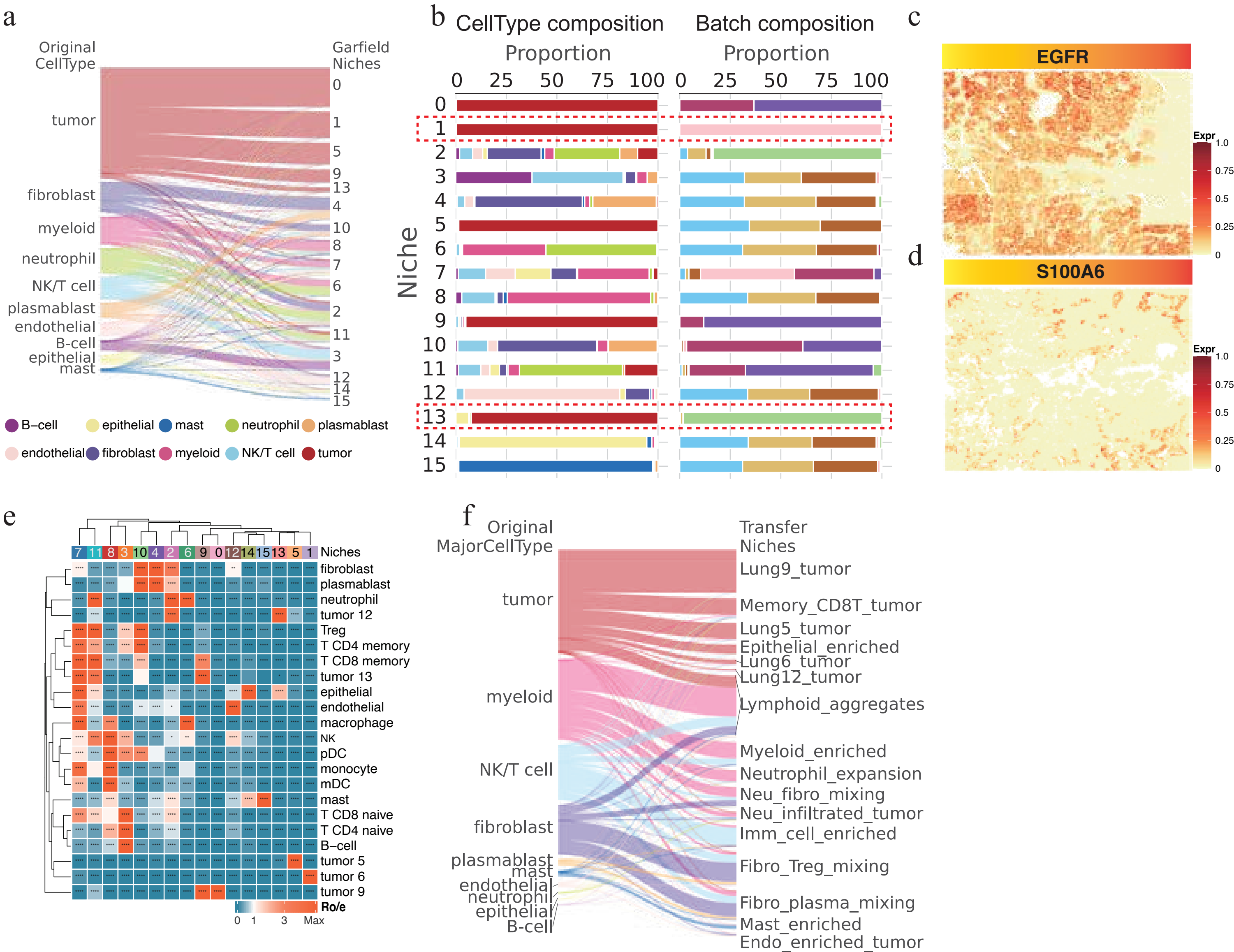

### Fig.S9

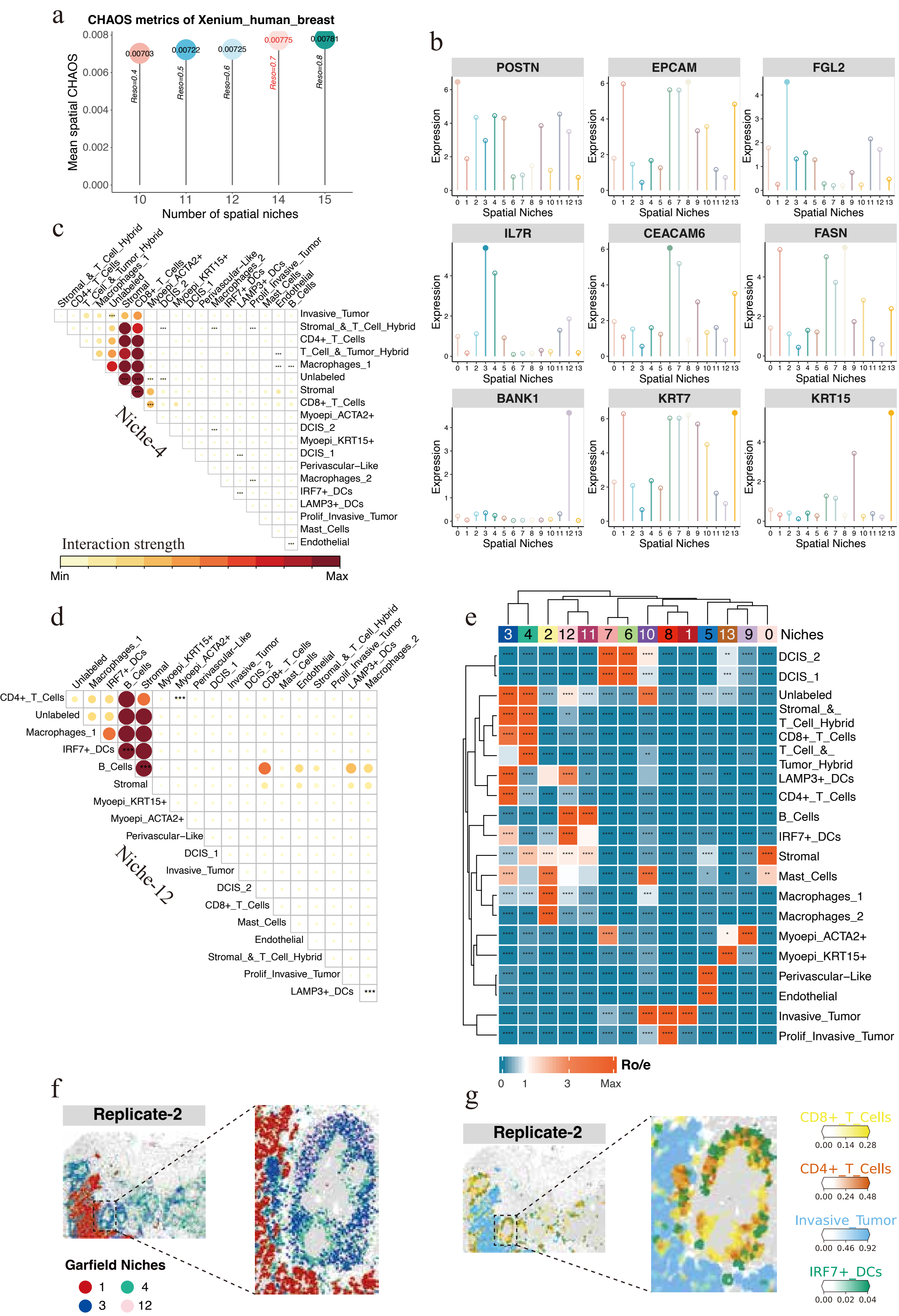
