## Supplementary material for "Graph-based Contrastive Learning Enables Unified Integration and Niche Transfer Across Single-Cell and Spatial Multi-Omics": Fig.S5

a

| Model | 4_run1 (0) | | | | | 8_run3 (1) | | | | | 12_run5 (2) | | | | | 16_run7 (3) | | | | | Aggregates | | | |
|---|---|---|---|---|---|---|---|---|---|---|---|---|---|---|---|---|---|---|---|---|---|---|---|---|
| | ARI | NMI | AMI | HOM | ASW | ARI | NMI | AMI | HOM | ASW | ARI | NMI | AMI | HOM | ASW | ARI | NMI | AMI | HOM | ASW | Overall Score (0) | Overall Score (1) | Overall Score (2) | Overall Score (3) |
| **Garfield** | 0.730 | 0.733 | 0.733 | 0.751 | 0.610 | 0.722 | 0.724 | 0.724 | 0.733 | 0.608 | 0.704 | 0.702 | 0.702 | 0.711 | 0.606 | 0.624 | 0.664 | 0.664 | 0.627 | 0.604 | 0.711 | 0.702 | 0.685 | 0.637 |
| **Cellcharter** | 0.554 | 0.598 | 0.598 | 0.600 | 0.554 | 0.560 | 0.592 | 0.592 | 0.596 | 0.559 | 0.527 | 0.577 | 0.577 | 0.580 | 0.561 | 0.564 | 0.592 | 0.592 | 0.590 | 0.563 | 0.581 | 0.580 | 0.564 | 0.580 |
| **Nichecompass** | 0.510 | 0.580 | 0.579 | 0.597 | 0.594 | 0.341 | 0.506 | 0.505 | 0.571 | 0.579 | 0.299 | 0.475 | 0.475 | 0.560 | 0.566 | 0.324 | 0.482 | 0.482 | 0.545 | 0.562 | 0.572 | 0.500 | 0.475 | 0.479 |
| **Graphst** | 0.527 | 0.554 | 0.553 | 0.541 | 0.507 | 0.432 | 0.530 | 0.529 | 0.572 | 0.508 | 0.406 | 0.519 | 0.518 | 0.546 | 0.506 | 0.498 | 0.553 | 0.553 | 0.552 | 0.508 | 0.536 | 0.514 | 0.499 | 0.533 |

b

| Model | 4_run2 (0) | | | | | 8_run4 (1) | | | | | 12_run6 (2) | | | | | 16_run8 (3) | | | | | Aggregates | | | |
|---|---|---|---|---|---|---|---|---|---|---|---|---|---|---|---|---|---|---|---|---|---|---|---|---|
| | ARI | NMI | AMI | HOM | ASW | ARI | NMI | AMI | HOM | ASW | ARI | NMI | AMI | HOM | ASW | ARI | NMI | AMI | HOM | ASW | Overall Score (0) | Overall Score (1) | Overall Score (2) | Overall Score (3) |
| **Garfield** | 0.749 | 0.734 | 0.734 | 0.751 | 0.610 | 0.711 | 0.706 | 0.706 | 0.736 | 0.605 | 0.685 | 0.691 | 0.691 | 0.684 | 0.607 | 0.692 | 0.695 | 0.695 | 0.715 | 0.604 | 0.716 | 0.693 | 0.672 | 0.680 |
| **Cellcharter** | 0.563 | 0.598 | 0.598 | 0.603 | 0.553 | 0.556 | 0.595 | 0.594 | 0.593 | 0.560 | 0.539 | 0.579 | 0.579 | 0.581 | 0.563 | 0.535 | 0.574 | 0.574 | 0.585 | 0.566 | 0.583 | 0.580 | 0.568 | 0.567 |
| **Nichecompass** | 0.437 | 0.560 | 0.560 | 0.600 | 0.594 | 0.340 | 0.505 | 0.505 | 0.570 | 0.579 | 0.323 | 0.479 | 0.479 | 0.537 | 0.568 | 0.342 | 0.483 | 0.483 | 0.521 | 0.561 | 0.550 | 0.500 | 0.477 | 0.478 |
| **Graphst** | 0.407 | 0.508 | 0.507 | 0.539 | 0.508 | 0.435 | 0.532 | 0.531 | 0.579 | 0.508 | 0.493 | 0.538 | 0.538 | 0.548 | 0.507 | 0.495 | 0.558 | 0.558 | 0.574 | 0.508 | 0.494 | 0.517 | 0.525 | 0.539 |
